# openPrimeR for multiplex amplification of highly diverse templates

**DOI:** 10.1101/847574

**Authors:** Christoph Kreer, Matthias Döring, Nathalie Lehnen, Meryem S. Ercanoglu, Lutz Gieselmann, Domnica Luca, Kanika Jain, Philipp Schommers, Nico Pfeifer, Florian Klein

## Abstract

To study the diversity of immune receptors and pathogens, multiplex PCR has become a central approach in research and diagnostics. However, insufficient primer design against highly diverse templates often prevents amplification and therefore limits the correct understanding of biological processes. Here, we present openPrimeR, an R-based tool for evaluating and designing multiplex PCR primers. openPrimeR provides a functional and intuitive interface and uses either a greedy algorithm or an integer linear program to compute the minimal set of primers that performs full target coverage. As proof of concept, we used openPrimeR to find optimal primer sets for the amplification of highly mutated immunoglobulins. Comprehensive analyses on specifically generated immunoglobulin variable gene segment libraries resulted in the composition of highly effective primer sets (oPR-IGHV, oPR-IGKV and oPR-IGLV) that demonstrated to be particularly suitable for the isolation of novel human antibodies.

## 1. Introduction

Multiprimer or multiplex polymerase chain reaction (mPCR) approaches are powerful tools to amplify target regions from multiple sites. They are of great importance for lymphocyte receptor profiling, pathogen identification, genotyping, and diagnostic pipelines. However, various targets exhibit a high degree of diversity, which challenges a robust amplification. As a result, sequence analysis is often compromised. For example, numerous viruses, such as the human immunodeficiency virus 1 (HIV-1) or the hepatitis C virus (HCV), possess error-prone replication machineries driving extensive sequence diversity. This can lead to a rapid development of escape variants under immunological or therapeutic pressure. In order to reliably detect escape from antiviral drugs, a comprehensive sequence analysis of virus variants is critical and of high relevance for a successful therapy (Beerenwinkel et al., 2001; Taylor et al., 2008; Doring et al., 2018; Bailey et al., 2019).

Other examples of highly diverse targets are B cell receptors (BCR). Here, mPCR primers need to bind the various V (variable) gene segments of the immunoglobulin loci (IGH, IGκ IGλ) that encode for the antibodies’ heavy and light chains (Wardemann et al., 2003; Tiller et al., 2008). Comprehensive analyses of the antibody repertoire can be exploited to elucidate the humoral immune response to vaccines or pathogens (Scheid et al., 2011; Muellenbeck et al., 2013 2013; Pappas et al., 2014; Galson et al., 2015; Rollenske et al., 2018) and are a prerequisite for the isolation of new monoclonal antibodies (mAbs). In fact, >50 new mAbs have been approved over the last years by the FDA/EMA to treat various diseases such as cancer, autoimmune or hematological disorders (Kaplon and Reichert, 2018). In this regard, novel HIV-1-directed broadly neutralizing antibodies (bNAbs) have recently been demonstrated to be promising candidates for HIV-1 prevention and therapy (Scheid et al., 2011; Klein et al., 2013; Caskey et al., 2015; Caskey et al., 2017; Bar-On et al., 2018; Gautam et al., 2018; Mendoza et al., 2018). bNAbs, however, pose a particular challenge to mPCR as they frequently contain a high burden of somatic hypermutation (SHM) as well as insertions or deletions (Zhou et al., 2010; Scheid et al., 2011; Klein et al., 2013). Importantly, forward primers binding to the signal peptide encoding leader (L) region of V genes have been demonstrated to favor the amplification of such highly mutated variable regions (Scheid et al., 2011).

These and other examples demonstrate the need for a comprehensive and reliable sequence analysis of highly variable targets. However, this requires efficient multiplex PCR primers that maximize the template coverage. A theoretical solution to this problem is the design of one individual primer for each target. However, the number of different primers that can be combined in mPCR is restricted. The pivotal task in mPCR primer design is thus to identify the minimal set of multiplex-compatible primers that covers all templates. This represents a so-called set cover problem (Feige, 1998; Alon et al., 2006) whose complexity rapidly increases with the number of templates. Primer design tools are typically programmed to filter individual primers for preferable properties (e.g. GC content, GC clamp, or melting temperature) and some also include a set cover optimization (Pesole et al., 1998; Wang et al., 2004; Jabado et al., 2006; Bashir et al., 2007; Gardner et al., 2009; Hysom et al., 2012) (Table 1). Of those, only PriMux (Hysom et al., 2012; Gardner et al., 2014), a command line software package, is currently available. However, PriMax only allows approximating a minimal set of primer pairs without adjusting the desired binding region. Therefore, it can not be used for the analysis of escape variants, the profiling of mutation patterns in antibodies, or recombinant expression of distinct protein fragments, which all require the explicit selection of the regions to be amplified.

**Table 1.**
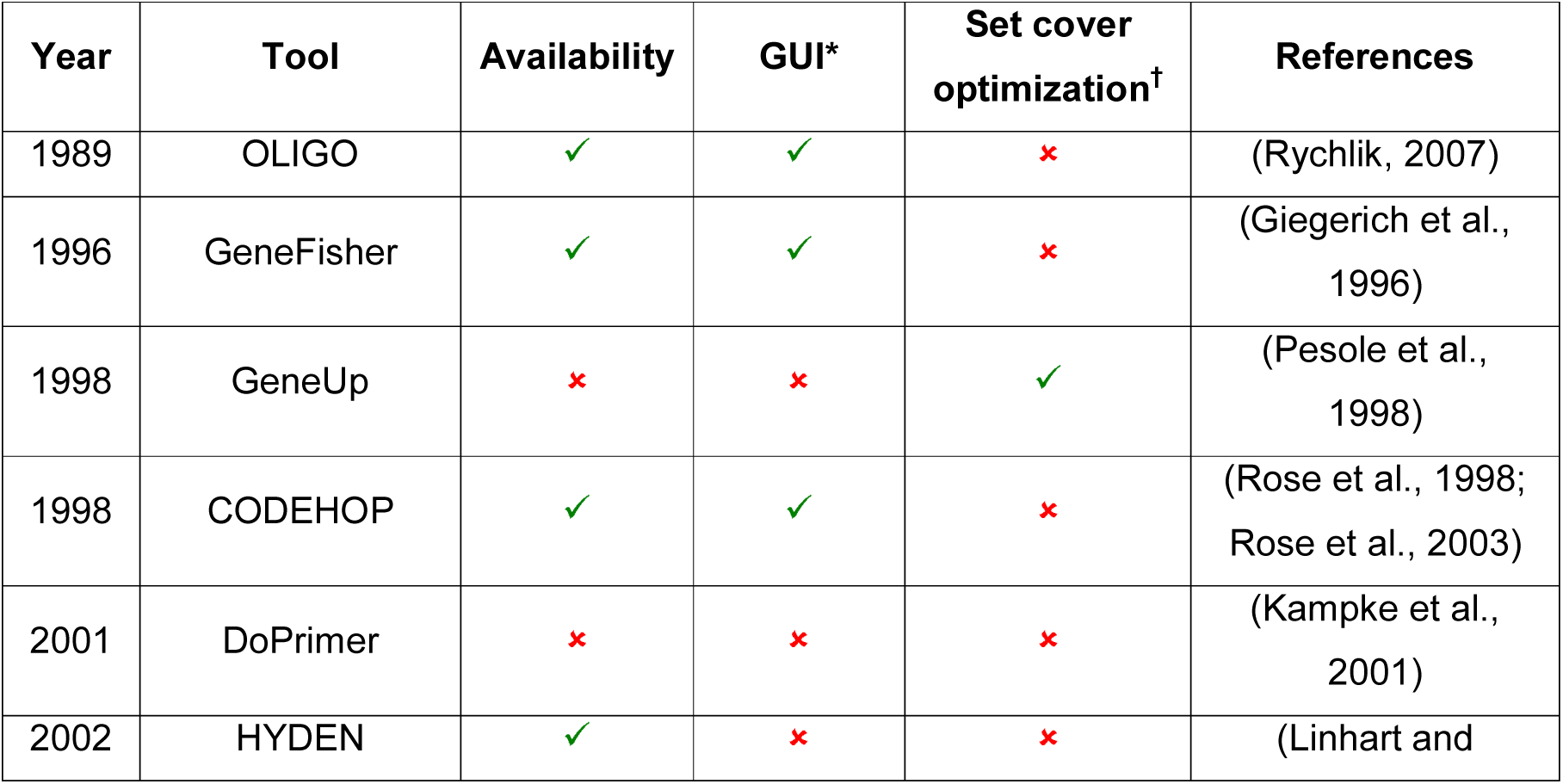
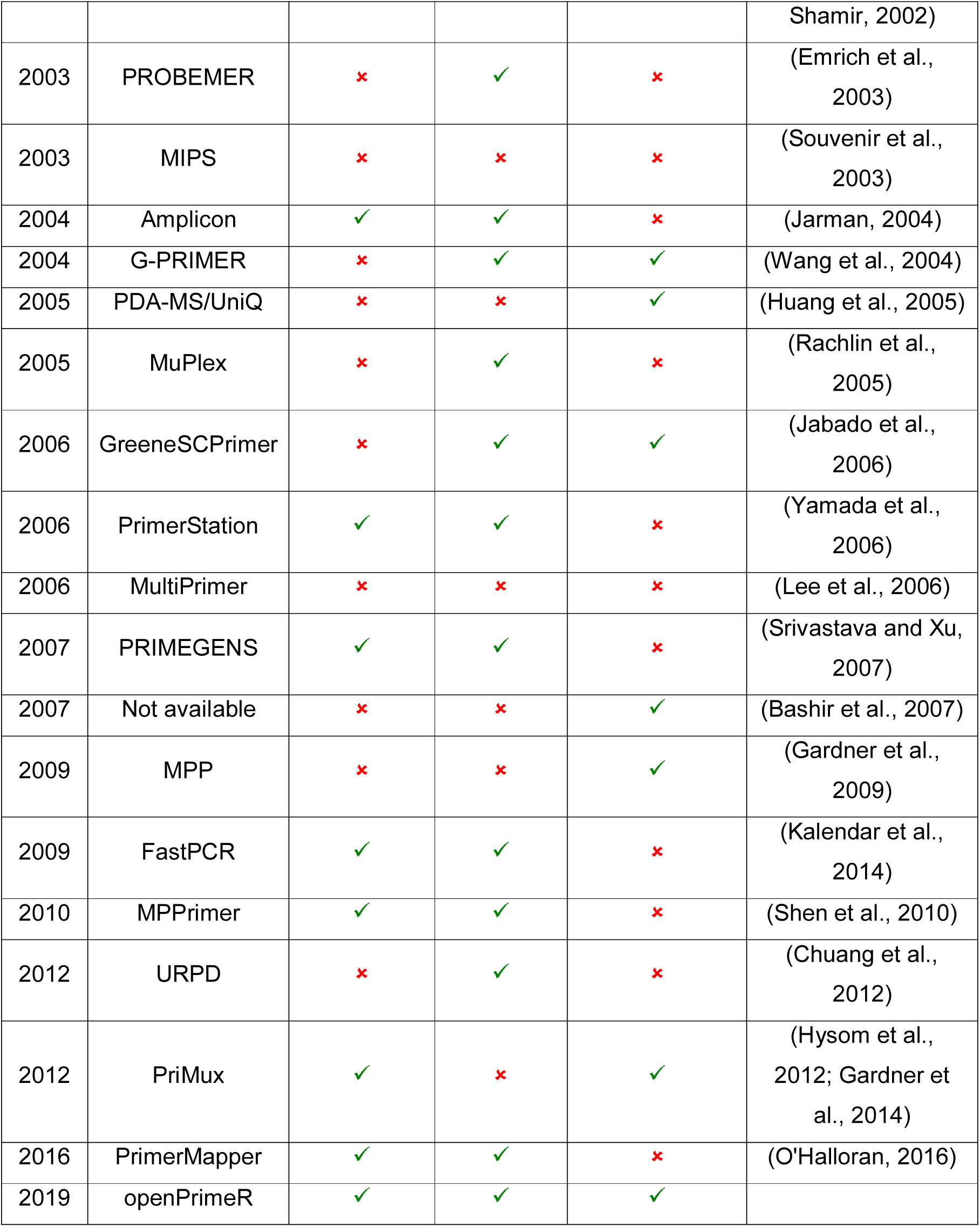
Approaches for multiplex primer design.

To address this problem, we developed openPrimeR (http://openprimer.mpi-inf.mpg.de/), a user-friendly program to i.) evaluate and ii.) design mPCR primers on a set of divers templates (Figure 1). openPrimeR, allows to exactly define the binding regions on multiple templates in parallel. It avoids primers that may form dimers or have other unfavorable physicochemical properties and it addresses the set cover problem by an integer linear program (ILP) or a greedy algorithm. In order to provide a versatile tool, we developed openPrimeR as an R package that allows other developers to include individual functionalities in their own primer design workflows. Moreover, we also equipped it with an intuitive graphical user interface and provide it as a docker container with all necessary dependencies. Combined with pre-defined default settings that have been validated extensively, openPrimeR can be used to design and to evaluate primers without computing skills or a deep understanding of mPCR chemistry. We used openPrimeR to evaluate published antibody-specific primer sets *in silico* and to design novel optimized primer sets that target the 5’ ends of the IGH, IGκ and IGλ V gene leader regions. We intensely validated openPrimeR and demonstrate the primers’ efficacy on specifically generated V gene libraries as well as on single B cells.

**Figure 1.**
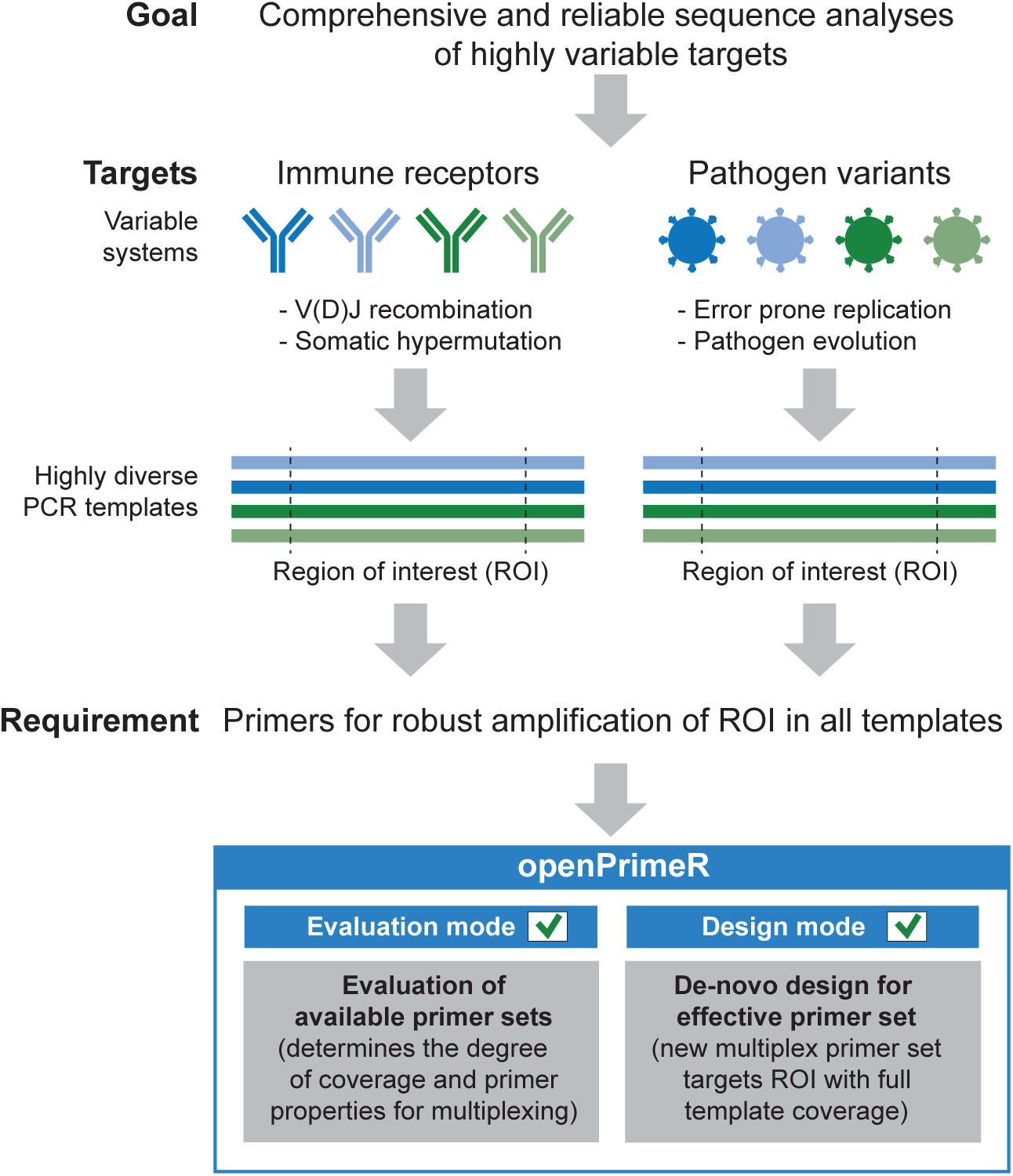
openPrimeR for evaluation and design of multiplex PCR primers against highly diverse templates. Sequence analysis of variable systems such as immune receptors or pathogens requires advanced primer sets for multiplex PCR amplification. openPrimeR supports both, the evaluation of existing primers as well as the *de-novo* design of a primer set that effectively amplifies a region of interest from a set of diverse templates.

## 2. Methods

### 2.1. Human samples

Blood samples from healthy and HIV-1-infected individuals were collected after written informed consent and in accordance with the requirements of the local Institutional Review Board (IRB; protocol number 16–110, University of Cologne, Cologne, Germany).

### 2.2. openPrimeR

openPrimeR is a multiplex primer evaluation and design tool that was developed in R. It provides a graphical user interface through a Shiny application and supports Linux, macOS, and Windows operating systems. For full functionality, openPrimeR requires installations of the additional programs MELTING (Le Novere, 2001), ViennaRNA (Lorenz et al., 2011), and OligoArrayAux (Markham and Zuker, 2008). openPrimeR provides two different modes: (1) evaluation, and (2) design. Both modes comprise a three-staged linear workflow consisting of (i) data input, (ii) settings configuration, and (iii) computation. In the input stage, templates are uploaded as FASTA/CSV files or chosen from integrated databases and the region of interest can be defined. In evaluation mode, primers can be uploaded as FASTA/CSV files or chosen from a selected set of previously published antibody-specific primer sets. In the configuration stage, up to 12 physicochemical property constraints, 5 coverage conditions, and 9 PCR settings can be specified (**Table S1**). Constraints are formulated in terms of permissible ranges of values. For example, a GC content constraint of [40%, 60%] indicates that primers should exhibit a GC content ranging from 40% to 60%. The final computation stage depends on the selected mode. Once computations are completed, results can be obtained directly as graphs or tables from the user interface or downloaded as CSV files for downstream analyses.

#### 2.2.1. Modes

In evaluation mode (1), computations comprise the determination of the physicochemical properties for each primer and all coverage events for each primer/template pair. The output includes tabular and graphical summaries of individual and overall coverage, binding positions, and violation of physicochemical property constraints. Primers violating active constraints can be removed from the set by a filtering function. Evaluation mode also provides a comparison function to directly compare multiple primer sets in parallel. In design mode (2), optimized primers are computed in three stages: (i) initialization, in which a set of candidate primers is generated, (ii) filtering, in which primers that violate active constraints are removed, and (iii) optimization, in which an instance of the set cover problem (SCP) is solved. Constraints may be relaxed during filtering and optimization to ensure that designed primers yield the user-defined target coverage ratio *CR* ∈ [0,1]. The extent to which constraints are relaxed is specified by the constraint limits. For example, given a constraint limit of [30%, 70%] for the GC clamp, the initial constraint on the GC content of [40%, 60%] could be relaxed to [30%, 70%].

#### 2.2.2. Templates

Virological and immunological templates were obtained from the Los Alamos National Laboratory HIV sequence database (Kuiken et al., 2003) (alignment of HIV subtype references, 2010) and the IMGT database (Lefranc et al., 1999) (retrieved April 2017), respectively. At the time of IMGT database access, 147 IGHV, 64 IGKV, and 36 IGLV templates with complete L1+V-Region sequences were available. In order to include at least one functional allele, functional V genes were supplemented with 5 IGHV, 2 IGKV, and 11 IGLV leader sequences retrieved from 5’RACE NGS data of naive B cells. In brief, PBMC from 8 healthy blood donors were enriched for CD19^+^ cells by MACS (Miltenyi Biotec) and 100,000 naive B cells (CD20^+^IgD^+^IgM^+^IgG^−^CD27^−^) were isolated by FACS for each individual. RNA was isolated (RNeasy Micro Kit; Qiagen) and cDNA prepared using the SMARTer RACE 5’/3’ Kit (Takara/Clontech). Samples were analyzed by 2×300 base pairs (bp) sequencing on an Illumina MiSeq at the Cologne Center for Genomics. Demultiplexed fastq-files were preprocessed using the pRESTO Toolkit (Vander Heiden et al., 2014) and self-written Python scripts. In short, forward and reverse reads were initially filtered for a minimum mean Phred score of 30 and sequencing lengths of at least 150 nucleotides (nt). Forward and reverse reads were assembled by requiring an overlap of at least 6 bp or concatenated if no overlap was found. Primer sequences were trimmed and reads were collapsed to extract unique sequences. Those sequences were analyzed by a stand-alone version of IgBLAST (Ye et al., 2013) and parsed with MakeDB.py from the Change-O Toolkit (Gupta et al., 2015). For leader sequence extraction, an in-frame ATG start codon was sought −51 to −66 nt upstream of framework region (FWR) 1. The sequence from the first start codon within this window up to the beginning of FWR1 was defined as the leader sequence. For each biological sample, leader sequences were then clustered according to their V gene allele. Intra-donor consensus sequences were generated for each V gene allele cluster within a donor by counting the occurrences of bases at each position and taking the most frequent as the consensus base (Schneider, 2002). A certainty score of each base position was calculated by dividing the count of an individual base by the total number of sequences within the cluster. Consensus leader sequences were built from groups of at least 10 unique reads of which none of the bases had a certainty score below 0.6. Leader sequences were used for database supplementation if they were found with 100% identity in at least 3 out of 8 different individuals.

#### 2.2.3. Initialization of Primers

For non-degenerate primers, candidate sequences are initialized as substrings from the template binding regions. Degenerate primers are initialized as described previously (Jabado et al., 2006). In brief, templates are aligned, subalignments corresponding to the desired primer length are extracted, pairwise sequence dissimilarities are determined, sequences are hierarchically clustered, and consensus sequences are formed along the constructed phylogenetic tree.

#### 2.2.4. Detection of Coverage Events

Potential primer binding regions are determined by considering all template substrings that match a primer with at most *n*_*mm*_ ≥ 0 mismatches. If a primer matches a template at multiple positions, the match with minimal *n*_*mm*_ is selected. Putative coverage events can be restricted in three ways. First, by considering the free energy of annealing and the presence of 3’ mismatches. Second, through the use of the thermodynamic model from DECIPHER (Wright et al., 2014) or a recently developed logistic regression model (Doring et al., 2019). Third, by removing coverage events inducing stop codons or amino acid mutations.

#### 2.2.5. Evaluation of Primer Properties

GC clamp, GC content, primer length, nucleotide runs and repeats are determined by text analysis. Melting temperature range, secondary structures, and self-dimerization are calculated with the external tools MELTING (Le Novere, 2001), ViennaRNA (Lorenz et al., 2011), and OligoArrayAux (Markham and Zuker, 2008), respectively.

#### 2.2.6. Optimization of Primers

Optimization of primers entails the selection of a minimal set of primers with a coverage ratio of least *CR* that fulfills the optimization constraints. There are two optimization constraints. One constraint on the maximal allowed melting temperature difference, *ΔT*_*m*_ ≥ 0, and one on the minimal allowed free energy of cross dimerization, 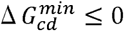. Compatible melting temperatures are ensured by forming primer subsets for individual melting temperature ranges [*T*_1_, *T*_2_] with *T*_2_ − *T*_1_ ≤ *ΔT*_*m*_ and solving the SCP for every subset. Once solutions for all primer subsets are obtained, the primer set with the maximal coverage at the smallest number of primers is chosen. Primer cross dimerization is prevented by constructing a binary dimerization matrix. Given a primer set *p* = {*p*_1_,*p*_2_,…,*p*_|*P*|_} and let Δ*G*(*p*_*i*_,*p*_*j*_) indicate the free energy of dimerization for primers *p*_*i*_ and *p*_*j*_. Then, the dimerization matrix *D* ∈ {0,1}^|*P*|×|*P*|^ is defined by

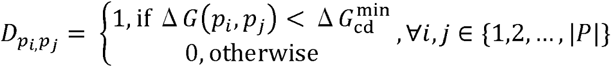

such that *D*_*pi,pj*_ =1 indicates that primers *p*_*i*_ and *p*_*j*_ dimerize according to 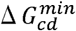. In order to solve the SCP instance, a greedy algorithm or an integer linear program (ILP) is used. The greedy algorithm selects a compatible primer with maximal coverage in every iteration. A candidate primer *p*_*i*_ is said to be compatible if *D*_*pi,pj*_ = 0 holds for all selected primers *p*_*j*_. Once primer *p*_*i*_ has been selected, the coverage of the remaining primers is updated by discounting the coverage events associated with *p*_*i*_. The algorithm terminates, if a coverage of *C* has been reached or when no compatible primers remain. The ILP for primer design requires the following definitions. Let *T* = {*t*_1_,*t*_2_,…,,*t*_|*T*|_} indicate the set of templates. The indicator vector *x* ∈ {0,1}^|*P*|^ shows whether *p*_*j*_ is included in the optimal set or not by defining it as

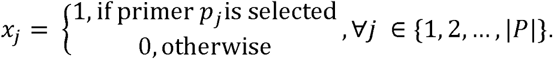

The coverage information is summarized in the coverage matrix *C* ∈ {0,1}^|*T*|×|*P*|^, which is defined by

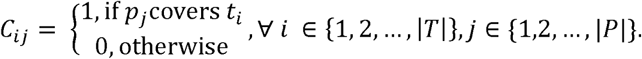

The ILP is given by

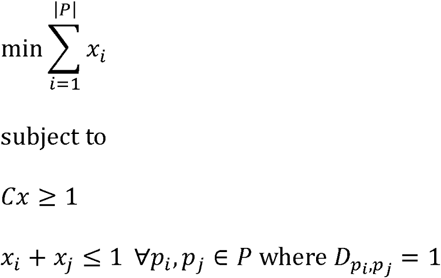

The target function ensures that the number of the selected set of primers is minimal. The first side constraint ensures that each template is covered by at least one primer and the second side constraint prevents cross-dimerizing primers. The ILP is solved using the exact branch-and-bound algorithm from lpsolve (Berkelaar et al., 2004).

### 2.3. Evaluation and comparison of published primer sets

For published primer sets, which were intended for nested PCR, we only considered forward primers of the first PCR for evaluation. This was based on the assumption that templates, which are not covered by the first PCR primers, will have a lower chance to be amplified during the second PCR by nested primers. If degenerate primers were reported they were translated to all underlying unambiguous derivatives for the analysis. 5’ overhangs from Set 3 (Kuppers et al., 1993) were removed to only analyze the complementary region of the primer. For the evaluation of primer sets we defined the following constraints: a GC content between 35 - 65%, 1 to 3 G or C nucleotides at the 3’ end (GC clamp), a maximum of 7 mismatches when none of them introduces a stop codon, a maximum run length of 4, a maximum repeat length of 4, a maximum deviation of 5°C between the lowest and the highest melting temperatures in a set, self- and cross-dimers with a dG cutoff of < −5 kcal/mol, a minimum coverage of at least 1 template for a primer. Binding events were predicted using the coverage model (Doring et al., 2019) with a false positive rate of 0.06. Primer binding regions were visually double-checked. When primers showed possible off-target binding within the same V gene, only the first event closest to the 5’ end was considered for the set binding region.

### 2.4. Gene library preparation for IGHV, IGKV, and IGLV

For generating the IGHV template library, PBMCs were isolated from 16 healthy individuals and enriched for CD19^+^ B cells (Miltenyi Biotec). Two times 2.5 × 10^5^ of naive B cells (CD20^+^IgM^+^IgD^+^IgG^−^) were sorted and subjected to RNA isolation (RNeasy Micro Kit; Qiagen). Following the generation of cDNA using 5’RACE (rapid amplification of cDNA-ends; SMARTer RACE 5’/3’ Kit; Takara/Clontech), PCR amplification was performed using a Q5 polymerase (NEB) and an C_H_1-specific reverse primer (5’ATGGAGTCGGGAAGGAAGTC’3 (Ozawa et al., 2006)). PCR products were analyzed by gel electrophoresis, purified (NuceloSpin Gel & PCR Clean-up Kit; Macherey-Nagel), and amplicons cloned into pCR4-TOPO (Invitrogen). For IGVK and IGVL libraries, the identical processing pipeline was performed on two additional pools of 8 and 10 individuals, respectively, with 2 × 10^5^ CD20^+^IgM^+^IgD^+^IgG^−^CD27^−^ sorted cells. In total 450, 688, and 546 colonies were analyzed by PCR for IGHV, IGKV, and IGLV, respectively and amplicons subjected to Sanger sequencing. Following IgBLAST (Ye et al., 2013) analysis all sequences with full-length annotated matches were further filtered to have a maximum of 6 nucleotide mismatches with no mismatches in the leader and the first 30 nucleotides of FWR1. Among 398 analyzed IGHV sequences, 10 IGHV genes could not be found and were thus synthesized (Eurofins genomics). All plasmids were adjusted to equal concentrations, aliquoted on hundred 96-well plates and stored at −20°C until used for the assays. An empty backbone plasmid was used as a negative control.

### 2.5. mPCR on V gene libraries

Multiprimer PCRs were performed in a volume of 25 µL using Taq Polymerase on 1 ng of template. Primers were multiplexed in an equimolar ratio with a final concentration of 200 nM primer mix together with 200 nM of an C_H_1 specific reverse primer (GGTTGGGGCGGATGCACTCC (Ippolito et al., 2012)). PCRs were performed with 94°C for 2 min, followed by 25 cycles 94°C for 30 sec, 57°C for 30 sec and 72°C for 55 sec, and a final step at 72°C for 5 min.

### 2.6. mPCR on single cell templates

PBMCs from 10 healthy donors were isolated by density gradient centrifugation (Ficoll– Paque; GE Healthcare) and enriched using human-CD19 coupled magnetic microbeads (Miltenyi biotech). Single naive (CD20^+^IgM^+^IgD^+^IgG^−^CD27^−^) and antigen-experienced (CD20^+^IgG^+^) B cells were sorted (Aria Fusion, BD) into 96-well plates and subjected to generate cDNA. PBMCs were also derived from a leukapheresis sample of an HIV-1-infected individual. In this case, B cells were enriched with the B cell isolation Kit II (Miltenyi biotec) and stained with a recombinant, GFP-tagged envelope trimer (Sliepen et al., 2015) to isolate HIV-1-reactive class-switched memory B cells (CD19^+^IgG^+^, see Figure S3 for gating strategies). Single cell analysis was performed as previously described (Tiller et al., 2008). In brief, single cell templates were lysed and reverse-transcribed using Superscript IV (Thermo Fisher) and random hexamer primers (Thermo Fisher). cDNA was subjected to PCR amplification using the conditions as indicated in **Table S4**. All PCR reactions were performed with PlatinumTaq (Thermo Fisher). As a PCR negative control, reverse transcription was either performed on empty wells (containing no single cells) or water was used instead of cDNA.

### 2.7. Determining PCR coverage

PCR products were supplemented with 6 x loading dye (Thermo Fisher) and visualized with SYBR Safe (Invitrogen) on a 2% agarose gel. In order to minimize biases in data analysis, all gel pictures were blinded by random coding and amplicon bands were independently labeled by five researchers. Labeling was decoded binary with 1 meaning any visible and definable signal at the expected height, and 0 meaning no signal at the expected height. The labels were combined and individual wells annotated as amplified at a cutoff score of three out of five positive ratings (**Figure S2, Figure S4, Figure S5, Figure S6, and Figure S7**).

### 2.8. Comparison of V gene identities

All positively scored wells from the oPR-IGHV PCRs on HIV-1-reactive B cells and all PCR products generated with Set 1 and/or Set 2 but not oPR-IGHV were analyzed by Sanger sequencing and annotated using IgBLAST (Ye et al., 2013). Sequences containing more than 15 bases with a Phred score below 16 or an average Phred score below 28 were excluded from further analyses yielding 179 high-quality and fully annotated heavy chain sequences. Plate and well position of V_H_ gene segments with < 70% germline V gene identity were identified and counted for the individual primer sets.

### 2.9. Quantification and statistical analysis

All statistical analyses were performed with Graphpad Prism (version 7.0b). All bar graphs depict mean values with standard deviations as error bars. P-values in Figure 3 were calculated using one-way ANOVA for matched data with Tukey test correction for multiple testing. The significance level was set at 0.05.

### 2.10. Data and software availability

#### 2.10.1. openPrimeR

openPrimeR is available in terms of two R packages that can be obtained via Bioconductor: (i) *openPrimeR* (https://bioconductor.org/packages/release/bioc/html/openPrimeR.html), which provides a programmatic interface, and (ii) *openPrimeRui* (https://bioconductor.org/packages/release/bioc/html/openPrimeRui.html), which provides a graphical user interface. The code for both projects is available via GitHub (https://github.com/matdoering/openPrimeR-User). A Docker container that includes all dependencies of openPrimeR is available at Docker Hub (https://hub.docker.com/r/mdoering88/openprimer/).

#### 2.10.2. Data availability

All data supporting the findings of this study are available within the paper and its supplementary information files. Extended IGHV, IGKV, and IGLV leader sequences are available via openPrimeR.

### 2.11. Reagents and Ressources

All reagents and resources used to conduct this work are listed in Table 2.

**Table 2.**
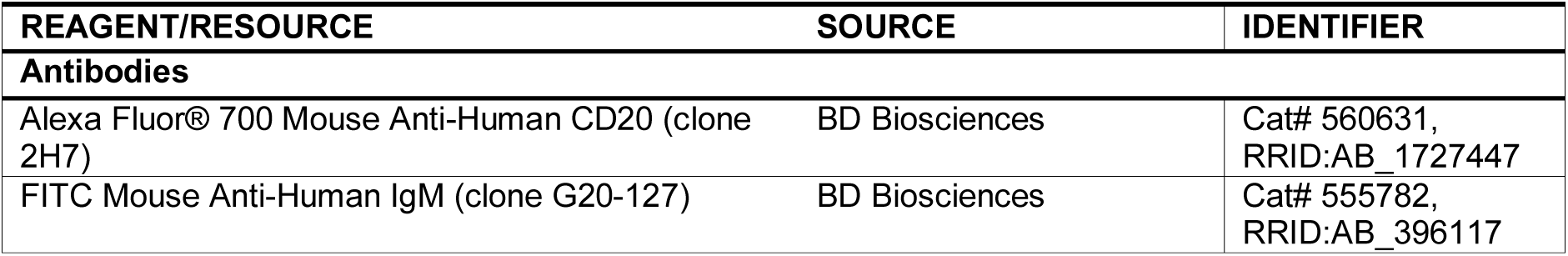
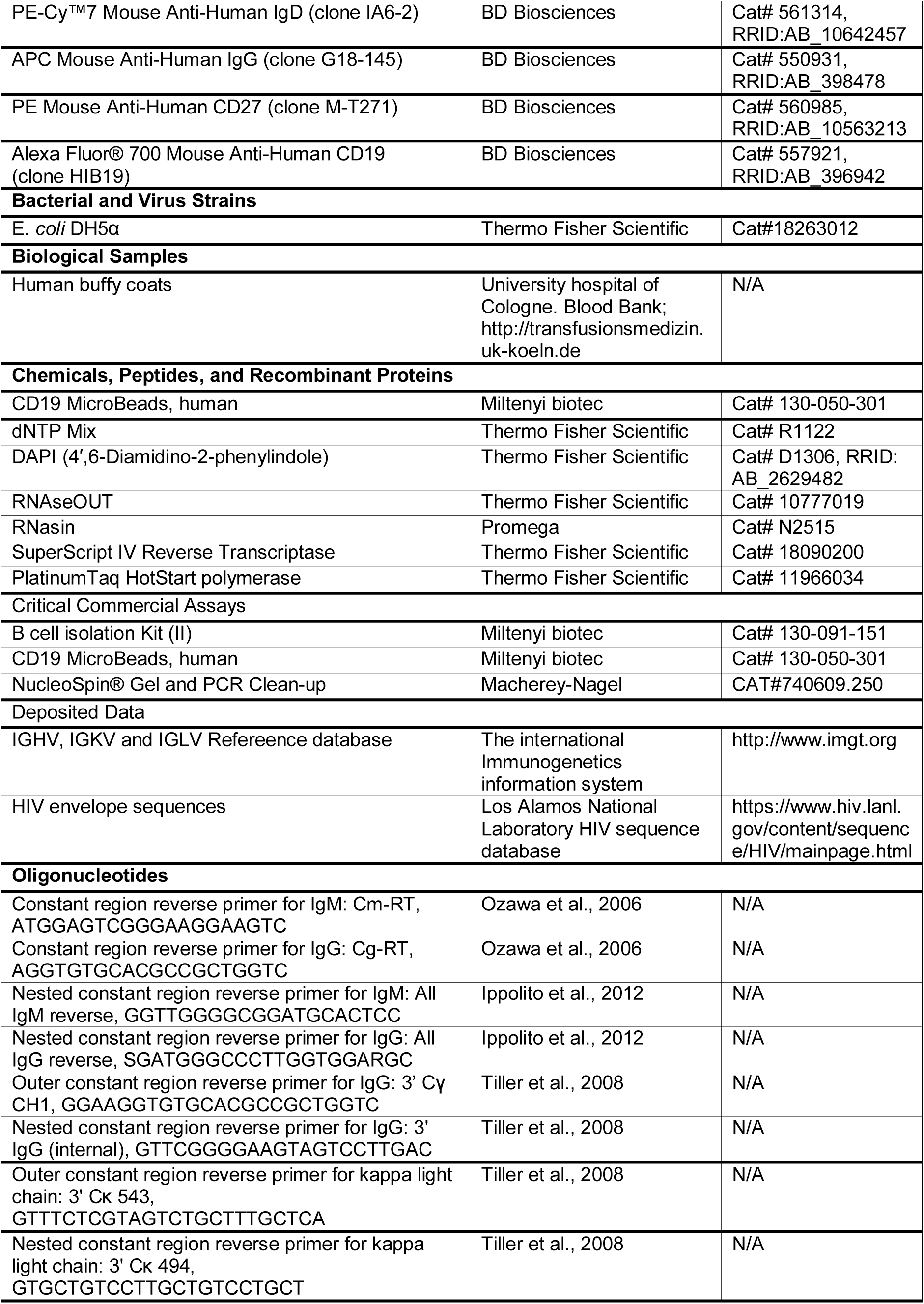
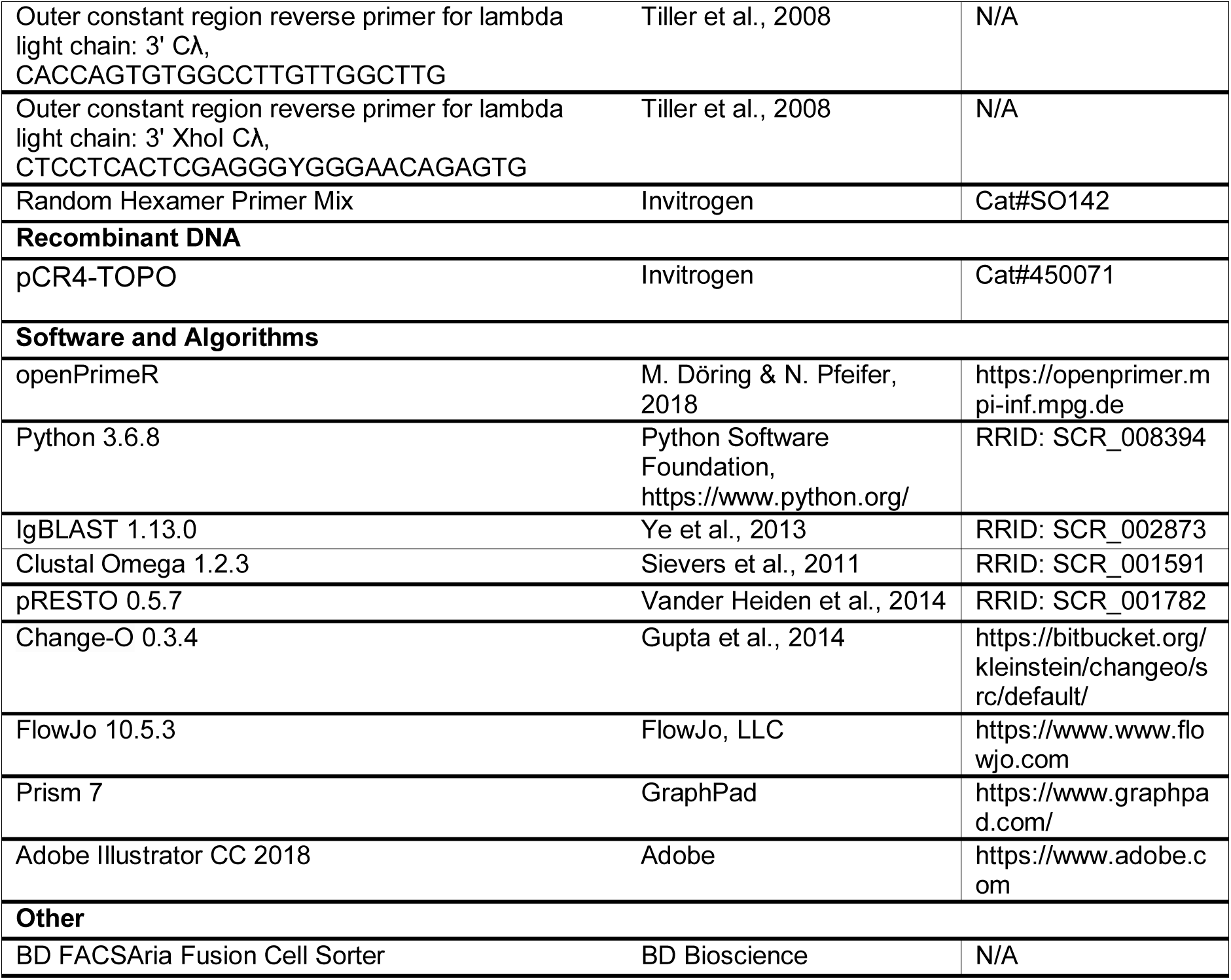
Reagents and resources.

## 3. Results

### 3.1. openPrimeR for evaluation and design of multiplex primer sets

openPrimeR is an R-based tool that includes modes for the evaluation and design of multiplex PCR primers targeting highly diverse templates such as immune receptors or fast evolving pathogens (Figure 1). It can be either accessed programmatically through R, or through the graphical user interface (GUI) of a Shiny application (**Figure S1**). The GUI is subdivided into an input part (left panel) that guides the user step-wise through the settings and an output part (right panel) that contains several tabs for selected information and output options. The user starts by uploading template sequences or choosing from integrated data, such as numerous sets of immunoglobulins (IMGT database (Lefranc et al., 1999)) or HIV reference sequence libraries (Los Alamos National Laboratory HIV sequence database (Kuiken et al., 2003)). The target regions can either be defined by uploading additional files containing the allowed binding regions for forward and reverse primers, or by defining them with slider ranges (most appropriate for sequences of identical length or with the same starting point).

In evaluation mode, primer sets are loaded into openPrimeR to determine their characteristics and to estimate their performance in a multiplex PCR experiment on the selected template sequences (Figure 2). In total, 12 physicochemical properties (e.g. melting temperature, GC content, GC clamp) and 5 coverage conditions (e.g. number of mismatches, introduction of stop codons, amino acid substitutions) can be considered (Figure 2, **Table S1**). In addition, it is possible to calculate the template coverage through an exact as well as an approximate string matching (with up to 20 mismatches) by using one out of three different models for the prediction of amplification events (Wright et al., 2014; Doring et al., 2019). As a result, tabular and graphical outputs offer a quick overview on whether the primer set covers the region of interest and the required multiplex properties.

**Figure 2.**
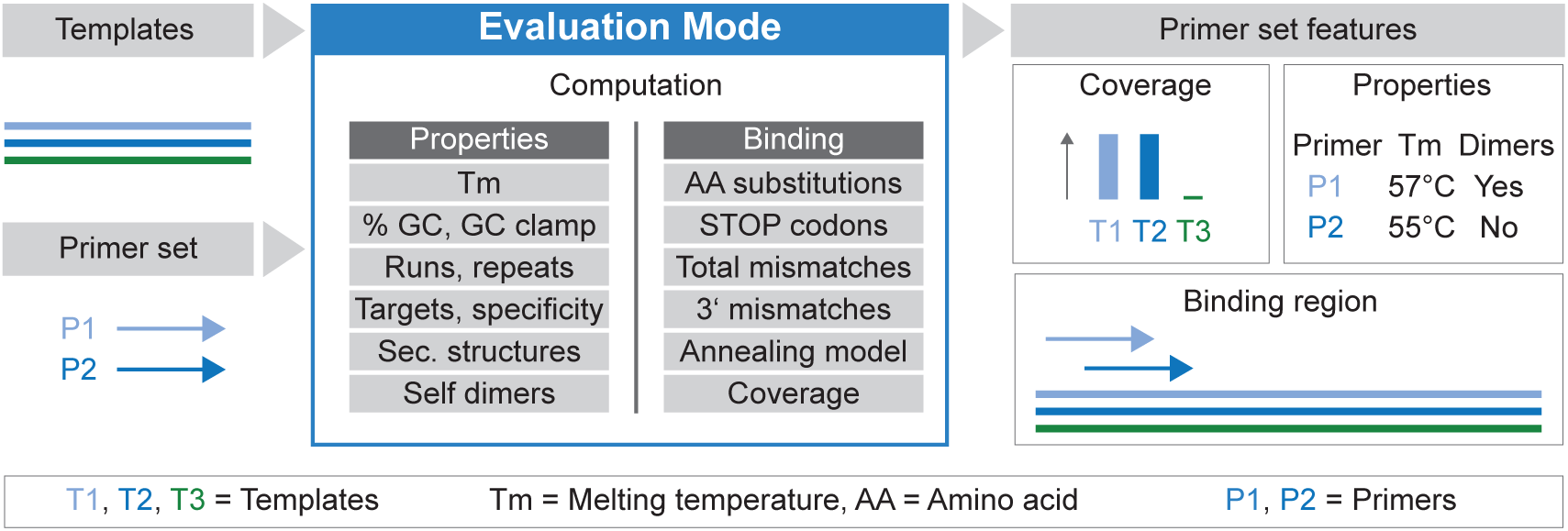
openPrimeR workflow for primer evaluation. Templates and primer sets are uploaded into openPrimeR. The software computes thermodynamic primer properties and predicts binding events for a selected region of interest. This allows a quick evaluation, whether a primer set will amplify the region of interest from all input sequences in a multiplex PCR. See also Table S1.

In the design mode, openPrimeR performs a stepwise primer generation process that results in the prediction of an optimal primer set (Figure 3). After defining the region of interest, openPrimeR determines all potential primer candidates within the selected range of lengths (initialization phase). If requested, primers are generated with a defined degree of degeneracy (see Methods section 2.2.3). Subsequently, based on the selected properties and conditions (**Table S1**), primers with unfavorable features will be removed from the initial set (filtering phase). During filtering, physicochemical constraints can be automatically relaxed to guarantee full target coverage. This abolishes the need to manually fine-tune constraints individually and start the process all over again. For example, adjusting the GC content to 40-60% might remove a good primer that covers 50% of the targets but has a GC content of 39%. Setting the lower GC content boundary to 35% will rescue this primer automatically, if the overall coverage drops below 100%. After filtering, a minimal set of primers with maximal coverage is selected using a greedy algorithm or an integer linear program (optimization phase). To ensure that selected primers can be multiplexed, the optimization procedure considers the maximal difference in primer melting temperatures and potential cross-dimerization events. If necessary, these multiplex constraints can be relaxed as the filtering constraints before (Figure 3, **Table S1**).

**Figure 3.**
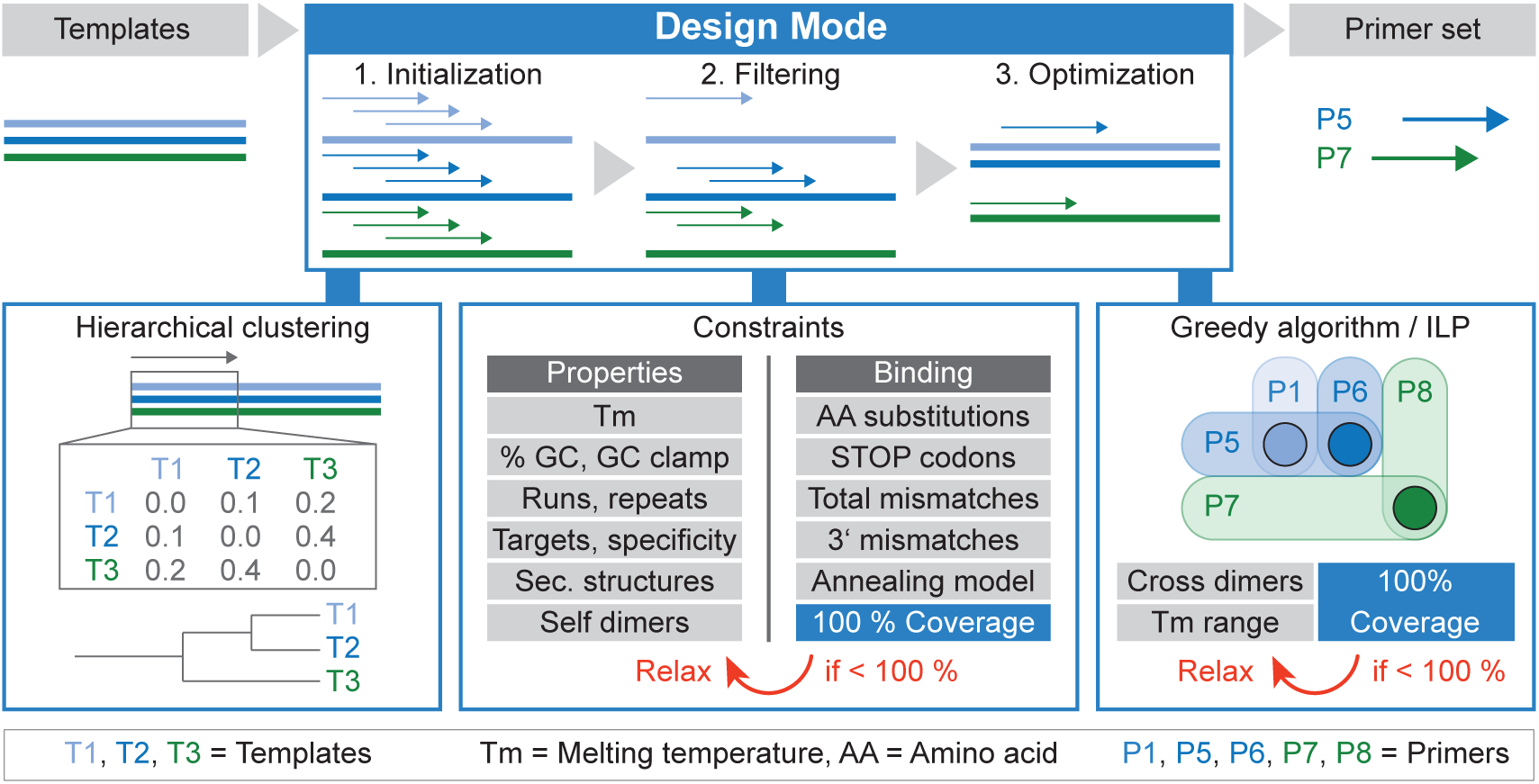
openPrimeR workflow for de novo primer design. Templates are uploaded into openPrimeR and a target region can be selected. An initial set of primers of specified length is either constructed by extracting template subsequences (non-degenerate primers) or by aligning and clustering of template subsequences (degenerate primers). Next, coverage is determined and primers are filtered according to user-defined constraints on their physicochemical properties. Finally, either a greedy algorithm or an integer linear program (ILP) finds an approximate (Greedy: P1, P6, P7) or exact (ILP: P5 and P7) minimal set of primers that covers all templates. If the coverage level is not achieved, constraints can automatically be relaxed to user-defined boundaries. See also Table S1.

### 3.2. openPrimeR reveals limitations for antibody sequence amplification

In the past, we and others have amplified B cell receptors (BCR) for antibody cloning and analysis (Kuppers et al., 1993; Sblattero and Bradbury, 1998; Wardemann et al., 2003; Tiller et al., 2008; Lim et al., 2010; Wu et al., 2010; Scheid et al., 2011; Ippolito et al., 2012; Klein et al., 2012; Murugan et al., 2015; Tan et al., 2016) using various sets of PCR primers. In order to evaluate their applicability in isolating highly mutated antibody sequences, we uploaded 6 representative primer sets (Kuppers et al., 1993; Tiller et al., 2008; Wu et al., 2010; Scheid et al., 2011; Ippolito et al., 2012; Tan et al., 2016) into openPrimeR. In evaluation mode their physicochemical properties and binding positions were computed for 152 functional IGHV gene segment alleles that were received from the IMGT database (Lefranc et al., 1999) and our own B cell repertoire sequencing data. Multiplex constraints for this evaluation comprised the GC content, the presence of a GC clamp, upper limits for runs and repeats, self- and cross-dimerization, and a limit for the melting temperature variation within a set. Binding events were considered positive with up to 7 mismatches (see Methods section 2.3.). The evaluation revealed that all tested primer sets carried some limitations (Figure 4), such as lacking the coverage of certain variable genes (Set 1, Set 5), performing only incomplete V gene segment amplification (Set 1, Set 2, Set 3, Set 4, Set 6), or requiring multiple reactions (Set 5). We conclude that no available primer set meets all criteria selected by openPrimeR for optimal amplification of antibody sequences.

**Figure 4.**
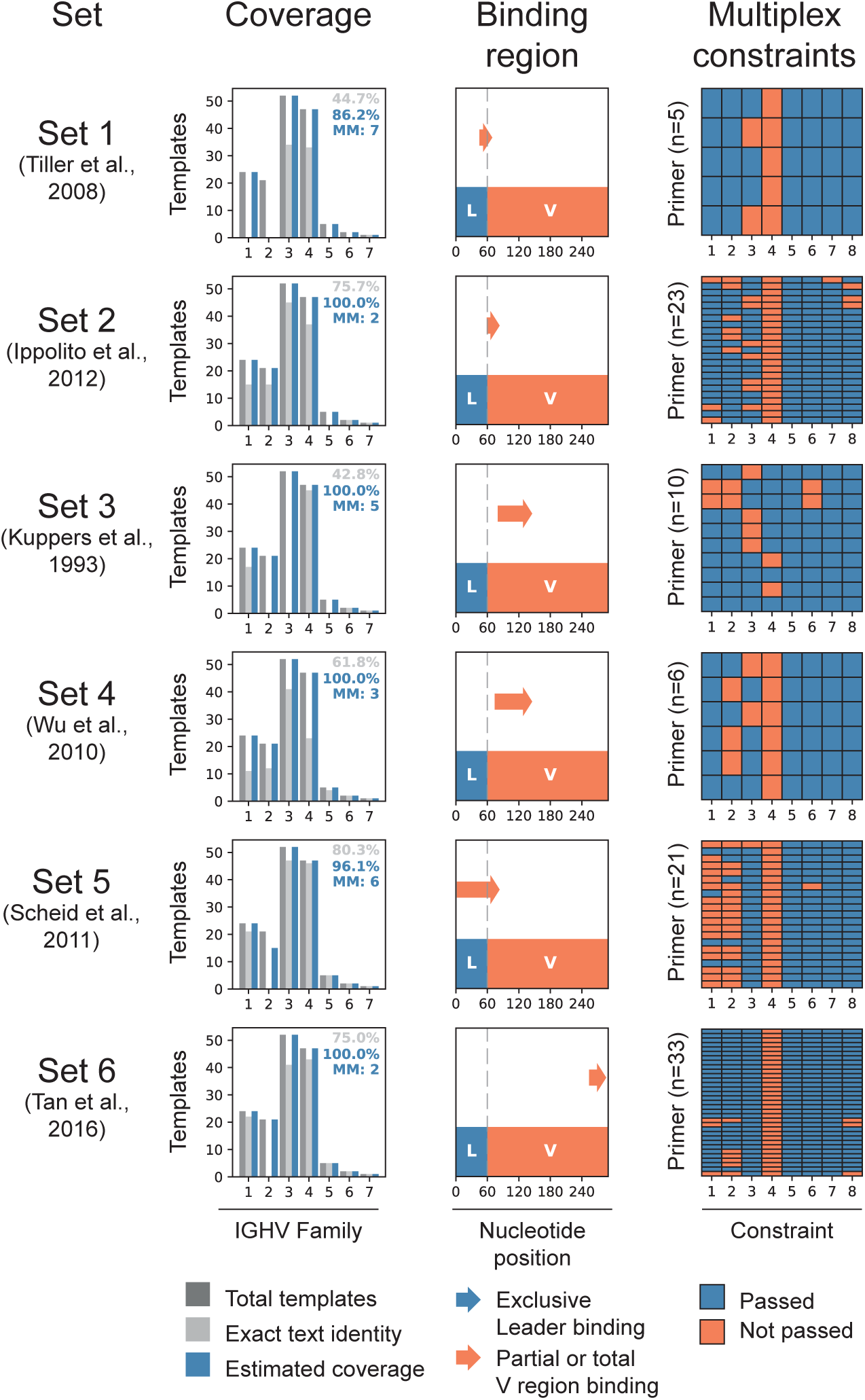
Evaluation of published primer sets using openPrimeR. Six representative primer sets that have been previously used to amplify V_H_ gene segments (Set 1 - 6) were analyzed with openPrimeR. Template coverage against 152 V_H_ gene segment alleles that can be subdivided into 7 families (dark grey bars) was determined by exact text identity (light grey) and maximum estimated coverage (blue). MM denotes the maximum number of 7 allowed mismatches that is necessary to reach the depicted estimated coverage. Binding regions are shown for all primers as a composite arrow that bridges the 5’ and 3’ ends of the outer most primers from the whole set. Each primer was tested for eight multiplex constraints (1-8: Cross dimers, GC clamp, GC ratio, melting temperature deviation, maximum number of nucleotide repeats, maximum number of nucleotide runs, coverage of at least 1 template, and self dimers). Constraints that were passed are colored in blue, those that were not passed are colored in red.

### 3.3. De novo openPrimeR-designed primer sets effectively amplify all V_H_, Vκ, and Vλ immunoglobulins

Since none of the published primer sets fulfilled our requirements, we set out to generate a novel optimized primer set for the amplification of human Ig heavy chain gene segments. In order to evaluate the quality of the openPrimeR design function, we produced a well-defined IGHV gene library containing one representative germline allele for all functional IGHV gene segments. The library was generated by template-switch-based 5’ rapid amplification of cDNA ends (5’RACE) of pooled naive B cells from 16 healthy individuals (Figure 5A). cDNA products were cloned and 450 plasmids were screened to collect full-length IGHV genes. Rare V_H_ gene segments that were not detected by this approach were sub-cloned from synthesized gene fragments resulting in a final library of 47 IGHV gene segments (**Table S2**). Next, we used openPrimeR to design IGHV primers that target the leader regions of all 152 functional IGHV gene alleles and tested them on the IGHV gene library. In an iterative process, we optimized starting conditions and filtering values (such as Gibbs energy cutoffs for primer-dimer prediction, number of allowed mismatches, etc.) that were eventually chosen as default settings for openPrimeR (Figure 5B and **Table S1**). The resulting set (oPR-IGHV, **Table S3**) contained 15 oligos with lengths between 20 to 28 nucleotides. These primers were predicted to bind exclusively to the leader region of all 152 sequences with a maximum of one mismatch, if not present at the very 3’ end (Figure 5C). We then compared the amplification of the IGHV library by oPR-IGHV with two of the previously published and frequently used antibody primer sets (Set 1 (Tiller et al., 2008) and Set 2 (Ippolito et al., 2012)) in five independent experiments. oPR-IGHV showed an overall coverage of 100% in four out of five experiments, whereas Sets 1 and 2 covered 95% and 97%, respectively (Figure 5D and **Figure S2**). Of note, Set 1 missed V genes IGHV2-5 and IGHV2-70 as predicted (Figure 4). V gene IGHV2-26, however, was amplified, suggesting that the set either introduced more than 7 mutations or stop codons by mismatch binding. We conclude that openPrimeR designs de novo primer sets that effectively amplify all V_H_ immunoglobulins.

**Figure 5.**
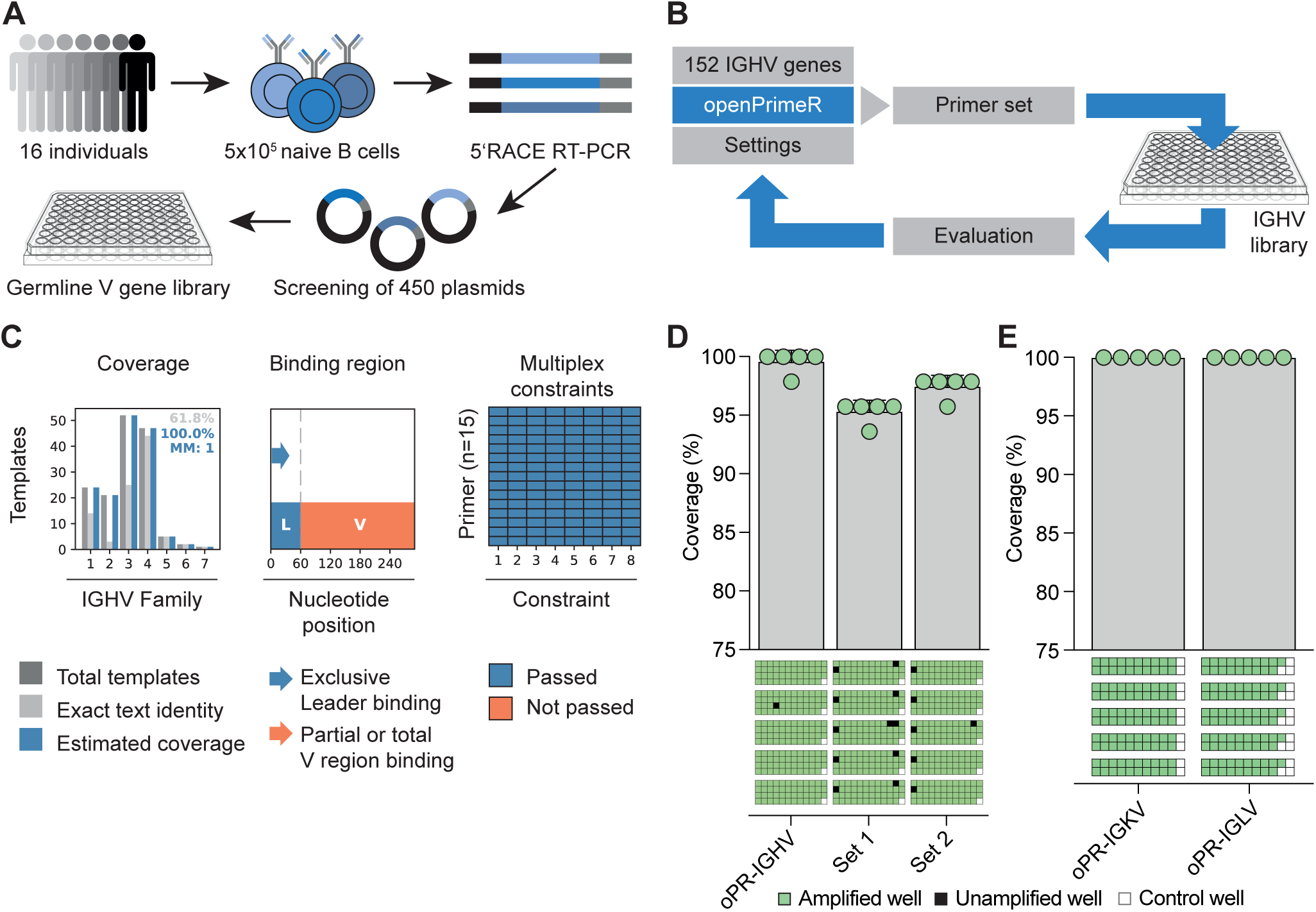
Validation of openPrimeR on specifically designed plasmid libraries. (**A**) In order to evaluate primer sets on defined templates, a germline IGHV gene library was established from pooled naive B cells of 16 healthy individuals. To this end, 5’RACE RT-PCR products were cloned into plasmids and one representative allele for each V gene was included into the library. (**B**) In an iterative process, primer sets were designed with openPrimeR and tested on the IGHV gene library to determine optimal default settings. (**C**) *In silico* re-evaluation of final IGHV primer set (oPR-IGHV) was performed in openPrimeR evaluation mode with the same settings as in Figure 4. (**D**) De novo oPR-IGHV primers were tested in five independent PCR experiments on the IGHV gene library in comparison to two published primer sets that have been used to isolate antibody sequences before. Bar graphs show mean coverage values over five experiments. (**E**) The same approach as in (A) was used to generate IGKV and IGLV gene libraries and IGKV/IGLV-specific primers were also designed with openPrimeR. Light chain primer sets (oPR-IGKV, oPR-IGLV) were tested in five independent PCR experiments on their respective gene library. Error bars in (**D**) and (**E**) depict standard deviations. Multiwell plates underneath the graphs represent individual experiments with green, black, and white squares indicating amplified, unamplified, and negative control wells, respectively. See also Figure S2, Table S1, Table S2 and Table S3.

To cross-validate the default settings on different templates, we used the same approach to design and test antibody light chain primers. All openPrimeR settings remained identical to the IGHV design process except for an increased stringency for 3’ mismatches (**Table S1**). As a result, openPrimeR predicted a set of 8 primers for IGKV and 15 primers for IGLV (oPR-IGKV and oPR-IGLV; **Table S3**). To test both primer sets we cloned and screened 1173 plasmids derived from B cell 5’RACE PCR products resulting in 20 and 19 different IGKV and IGLV genes, respectively (**Table S2**). Although these 39 genes represent only 53% of the 41 IGKV and 33 IGLV genes listed in the IMGT database, they accounted for over 90% of all light chain V gene segments that we found within the B cell receptor repertoires of 8 healthy individuals (see Method section 2.4). Both primer sets covered 100% of their respective Vκ/λ gene library (Figure 5E and **Figure S2**). Taken together, the results on IGHV, IGKV, and IGLV libraries demonstrate the capability of openPrimeR to select minimal sets of primers (oPR-IGHV oPR-IGKV, oPR-IGLV) that effectively amplify all intended target templates in multiplex PCRs.

### 3.4. openPrimeR-derived primers are superior in isolating highly mutated antibody sequences

After validating openPrimeR on antibody sequence libraries, we investigated oPR-IGHV amplification of the immunoglobulin locus from single B cells with different levels of somatic hypermutation. To this end, we isolated single naive B cells as well as antigen-experienced IgG^+^ memory B cells from healthy individuals expected to carry non/low and moderate levels of mutations, respectively. In addition, we used HIV-1-reactive memory B cells from an individual with broad HIV-1-reactive serum activity as a source of highly mutated antibody sequence templates (see **Figure S3** for sorting strategies). From a total of 841 sorted B cells, cDNA was generated and equally subjected to PCR reactions using primer sets 1, 2, and oPR-IGHV. In order to estimate successful amplification, PCR results were visualized by agarose gel electrophoresis, blinded, and evaluated by five independent investigators for the presence or absence of a PCR product at the correct position (**Figure S4, Figure S5, and Figure S6**). Among all three primer sets we did not detect significant differences in amplifying sequences from naive or IgG^+^ memory B cells from healthy individuals (Figure 6A, **Figure S4 and Figure S5**). However, the overall coverage of HIV-1-reactive B cells was substantially higher for oPR-IGHV (Figure 6A and **Figure S6**), yielding 21 quality-controlled heavy chain sequences (12%) that could not be identified with either Set 1 or Set 2. In order to investigate the influence of SHM on the coverage of the different sets, we determined the germline V_H_ gene identity of all oPR-IGHV-derived PCR products. All three primer sets had a comparable capacity to amplify mutated sequences down to 70% germline identity (Figure 6B). However, amplification of sequences with < 70% V gene identity was significantly more effective for oPR-IGHV than for Set 1 (p = 0.0025) and Set 2 (p = 0.0017) (Figure 6B). In addition, we detected germline reversion and primer-introduced mutations in the framework region 1 for sequences that were captured by Set 1 or Set 2 (data not shown). We conclude that oPR-IGHV is superior in amplifying highly mutated antibody sequences.

**Figure 6.**
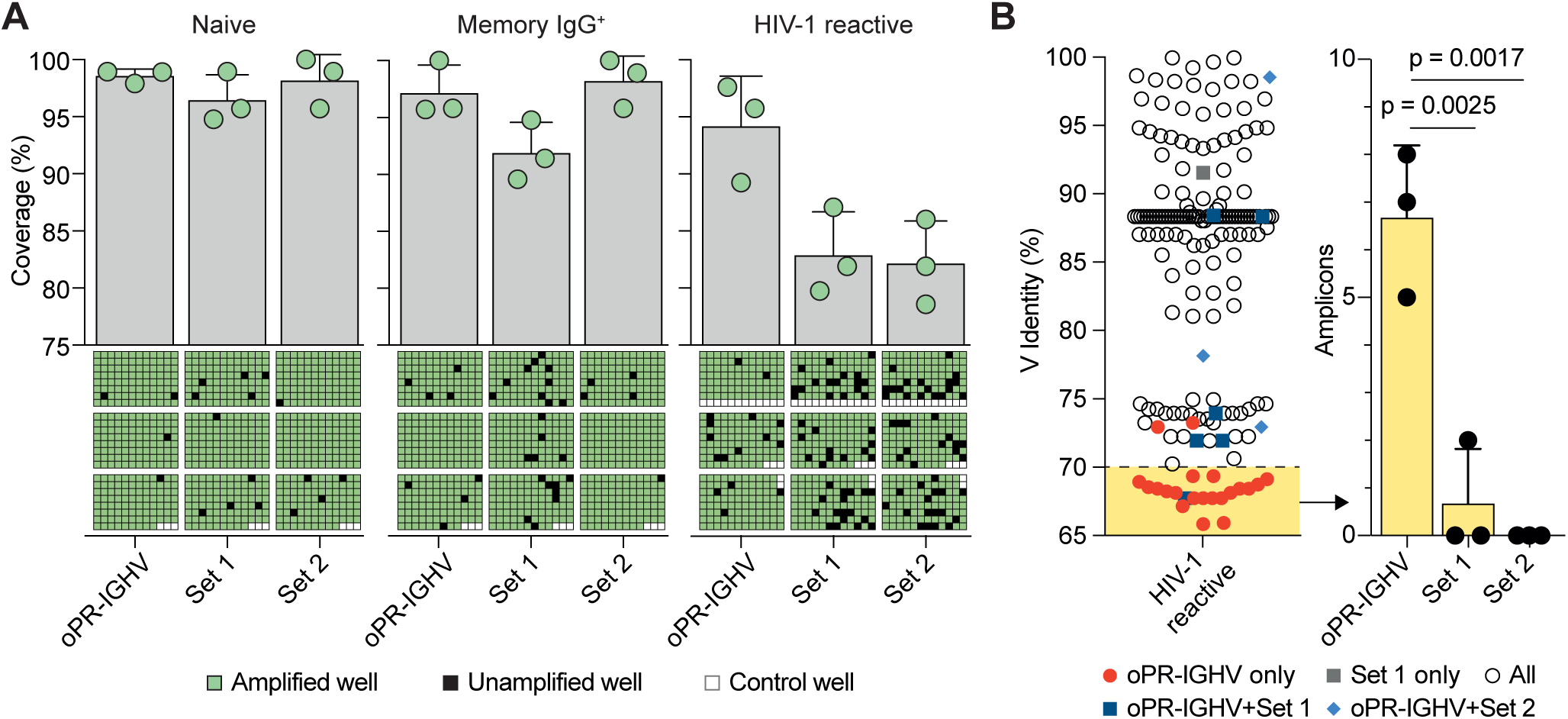
High performance of oPR-IGHV on amplifying diverse and somatically mutated antibody genes. (**A**) Successful amplification of BCR heavy chain sequences from single naive, memory IgG^+^, or HIV-1-reactive B cells performed by oPR-IGHV, Set 1, and Set 2, respectively. Individual experiments (n = 3 per B cell subset) are shown as schematic multiwell plates (See also Figure S4, Figure S5, and Figure S6). Amplified, unamplified, and negative control wells are depicted in green, black, and white, respectively. (**B**) Left panel: Heavy chain V gene identities from amplified HIV-1-reactive B cell receptors shown in (A). Red circles depict heavy chains that have only been amplified by oPR-IGHV primers (12% of all PCR products), blue symbols indicate heavy chains amplified by oPR-IGHV as well as one of the other sets. Right panel: Number of amplification events per cDNA run that had a V gene identity of < 70% for the three primer sets oPR-IGHV, Set 1, and Set 2. Bar graphs depict mean values and show standard deviations as error bars. One-way analysis of variance (ANOVA) for matched data was performed on the amplification events with < 70% V gene identity. P-values (Tukey post-hoc test) for pairwise comparisons are displayed, if they are below a significance level of 0.05.

Finally, corresponding light chains were amplified using oPR-IGKV and oPR-IGLV primer sets with a success rate of 98% (**Figure S7**). Of note, following cloning and production of antibodies, we were able to identify numerous broad and highly potent HIV-1 neutralizing antibodies with V gene identities of less than 70% (Schommers et al., manuscript in preparation). These antibodies were not detected using primer sets 1 or 2. We therefore conclude that the primer sets generated by openPrimeR are of great value to precisely identify antibody sequences independent of their mutational status.

## 4. Discussion

mPCR is a fundamental technique that requires advanced primer design. However, current design tools lack functionality for a set cover optimization (Giegerich et al., 1996; Rose et al., 1998; Kampke et al., 2001; Linhart and Shamir, 2002; Emrich et al., 2003; Souvenir et al., 2003; Jarman, 2004; Rachlin et al., 2005; Lee et al., 2006; Yamada et al., 2006; Rychlik, 2007; Srivastava and Xu, 2007; Shen et al., 2010; Chuang et al., 2012; Kalendar et al., 2014; O’Halloran, 2016), are not freely available (Pesole et al., 1998; Kampke et al., 2001; Emrich et al., 2003; Souvenir et al., 2003; Wang et al., 2004; Rachlin et al., 2005; Jabado et al., 2006; Lee et al., 2006; Bashir et al., 2007; Gardner et al., 2009; Chuang et al., 2012), or do not provide a graphical user interface for an intuitive workflow and data evaluation (Pesole et al., 1998; Kampke et al., 2001; Linhart and Shamir, 2002; Souvenir et al., 2003; Lee et al., 2006; Bashir et al., 2007; Gardner et al., 2009; Hysom et al., 2012). To overcome these limitations, we developed and experimentally tested openPrimeR, a user-friendly mPCR primer tool that can be used for i.) evaluating existing and ii.) designing novel and highly effective mPCR primer sets. Evaluation with openPrimeR can be used to quickly predict the performance of established primers *in silico* and thereby keep track of the coverage for large libraries of templates. In addition, predictions can be used to compare the performance of different primer sets on target template sequences. The design mode of openPrimeR is highly flexible and allows adjusting a large number of primer properties. However, by experimental validation, we identified reasonable default settings to assist in primer design even without a deep understanding of mPCR primer requirements. Importantly, the optimization algorithms in design mode automatically select the minimal number of primers that fulfill all mPCR requirements. This abolishes the need for tedious and cost-intensive experimental evaluation of possible primer combinations.

As a proof of concept, we applied openPrimeR to evaluate and design human antibody-specific primer sets. Various primer sets have successfully been used in single B cell cloning approaches to decipher B cell responses and to isolate monoclonal antibodies (Wardemann et al., 2003; Tiller et al., 2008; Scheid et al., 2011; Ippolito et al., 2012; DeKosky et al., 2013). However, evaluation with openPrimeR revealed certain limitations of existing primer sets. These include the lack of covering all currently known V genes (Tiller et al., 2008; Scheid et al., 2011), incomplete V gene segment amplification (Kuppers et al., 1993; Tiller et al., 2008; Wu et al., 2010; Ippolito et al., 2012; Sun et al., 2012; Tan et al., 2016), or elevated costs because of the requirement for several polymerase chain reactions (Scheid et al., 2011). We thus used openPrimeR to design IGHV, IGKV, and IGLV primer sets that facilitate efficient amplification of highly mutated heavy and light chain sequences and could demonstrate their efficiency on gene libraries and single cells. These novel sets have contributed to isolating broad and potent HIV-1 neutralizing antibodies in our own lab (unpublished data) and will be of great value for the precise amplification of antibody sequences independent on the level of somatic mutations.

We expect that openPrimeR will be of great value for all researchers that require successful and reliable amplification of diverse targets. openPrimeR includes 436 human IGHV, IGKV, and IGLV templates, 77 published antibody-specific primer sets, and is available at Bioconductor, as a Docker container, or from a GitHub repository (see Methods section 2.10.1.).

### 4.1. Limitations

openPrimeR is subject to the following limitations. When the optimization problem is solved using the greedy algorithm, the smallest possible primer set may not be found. Since the worst-case runtime of the ILP-formulation is exponential, primer design may not be feasible for large numbers of templates when this optimization strategy is used. Moreover, the overall runtime depends on the user-provided primer design settings (e.g. number of considered constraints, approach for coverage estimation). Runtime increases when the constraints on the physicochemical properties of primers have to be relaxed. In addition, openPrimeR was initially designed to assist in the primer design against immunological or virological targets, which have known regions of hypervariability. The tool therefore allows to set primer binding regions outside of mutational hotspots based on prior knowledge. If sequences for all expected mutation variants are available, openPrimeR could in principle be used to design degenerate primers within such mutational hotspots. For future releases, it could be an option to include interrogating template sequences for defined mutational hotspots (such as WRC/GYW for the activation induces deaminase (AID)) and automatically skip regions with such motifs.

## Acknowledgements

We would like to thank Thomas Lengauer and Henning Gruell for helpful comments on the manuscript and all members of the Klein Lab for helpful discussion. For testing openPrimeR, we thank Dilip Ariyur Durai, Sivarajan Karunanithi, Markus List, Fabian Müller, Sarvesh Nikumbh, Michael Scherer, and Florian Schmidt. We further thank Peter Nürnberg, Janine Altmüller and Christian Becker for NGS support. The work was supported by the Heisenberg-Program of the DFG (KL2389/2-1) (F.K.), the European Research Council (ERC-StG639961), the German Center for Infection Research (DZIF) (P.S., F.K.), and the German Research Foundation (CRC 1279, CRC 1310) (F.K., C.K., M.S.E.).

## Author contributions

CK and FK initiated the project and designed all experiments. MD developed openPrimeR. CK, NL, DL and LG performed experiments. CK, MD, LG, NL, MSE, and FK evaluated the data. PS, MSE and KJ acquired blood samples and performed FACS sorting. NP supervised the development of openPrimeR. CK, MD, and FK wrote the manuscript. All authors discussed the results and reviewed the manuscript.

## SUPPLEMENTAL INFORMATION

### Supplementary Figures

**Figure S1. openPrimeR graphical user interface.** A Shiny application can be used for interactive usage of openPrimeR. The left panel contains the input interfaces and guides the user step-wise through the evaluation or design mode. The right panel offers several tabs for output display.

**Figure S2. Visualization of PCR products for IGHV, IGKV, and IGLV gene libraries by gelelectrophoresis.** IGHV library was amplified with oPR-IGHV, Set 1, and Set 2. IGKV and IGLV gene libraries were amplified with oPR-IGKV and oPR-IGLV, respectively. Green, black, and white squares indicate amplified, unamplified, and control wells (not counted). Note: IGHV template in D4 turned out to have a deletion in the binding region of oPR-IGHV primer 9.

**Figure S3. Gating strategies for single cell sorting.** (**A**) Gating strategy for naive and antigen-experienced B cells. (**B**) Gating strategy for HIV-1-specific B cells.

**Figure S4. Visualization of PCR products from naive B cell cDNA with oPR-IGHV, Set 1, and Set 2.** Green, black, and white squares indicate amplified, unamplified, and control wells (not counted), respectively.

**Figure S5. Visualization of PCR products from IgG+ memory B cell cDNA with oPR-IGHV, Set 1, and Set 2.** Green, black, and white squares indicate amplified, unamplified, and control wells, respectively. Note: Controls and grey squares were not taken into account for coverage determination.

**Figure S6. Visualization of PCR products from HIV-1_BG505.SOSIP_-reactive B cell cDNA with oPR-IGHV, Set 1, and Set 2.** Green, black, and white squares indicate amplified, unamplified, and control wells (not counted), respectively.

**Figure S7. PCR results from HIV-1_BG505.SOSIP_-reactive B cell cDNA with oPR-IGKV and oPR-IGLV.** Light chain coverage summary. Kappa and lambda light chain specific PCRs were performed on HIV-1_BG505.SOSIP_-reactive B cells with oPR-IGKV and oPR-IGLV primer sets. Green, black, and white boxes depict amplified, unamplified, and control wells (not counted), respectively. Percentages indicate mean coverage ± standard deviation.

### Supplementary Tables

**Table S1. openPrimeR options, default settings and settings used for primer design on IGHV, IGKV, and IGLV**

**Table S2. IGH, IGK, and IGL V gene alleles used in standard libraries**

**Table S3. openPrimeR-derived optimized primer sets oPR-IGHV, oPR-IGKV, and oPR-IGLV for amplification of antibody heavy and light chains**

**Table S4. Primer combinations and thermocycling conditions for multiplex PCRs**

## References

Alon, N., Moshkovitz, D. and Safra, S., 2006, Algorithmic construction of sets for k-restrictions. ACM Trans. Algorithms 2, 153–177.

Bailey, J.R., Barnes, E. and Cox, A.L., 2019, Approaches, Progress, and Challenges to Hepatitis C Vaccine Development. Gastroenterology 156, 418–430.

Bar-On, Y., Gruell, H., Schoofs, T., Pai, J.A., Nogueira, L., Butler, A.L., Millard, K., Lehmann, C., Suarez, I., Oliveira, T.Y., Karagounis, T., Cohen, Y.Z., Wyen, C., Scholten, S., Handl, L., Belblidia, S., Dizon, J.P., Vehreschild, J.J., Witmer-Pack, M., Shimeliovich, I., Jain, K., Fiddike, K., Seaton, K.E., Yates, N.L., Horowitz, J., Gulick, R.M., Pfeifer, N., Tomaras, G.D., Seaman, M.S., Fatkenheuer, G., Caskey, M., Klein, F. and Nussenzweig, M.C., 2018, Safety and antiviral activity of combination HIV-1 broadly neutralizing antibodies in viremic individuals. Nat Med 24, 1701–1707.

Bashir, A., Liu, Y.T., Raphael, B.J., Carson, D. and Bafna, V., 2007, Optimization of primer design for the detection of variable genomic lesions in cancer. Bioinformatics 23, 2807–15.

Beerenwinkel, N., Lengauer, T., Selbig, J., Schmidt, B., Walter, H., Korn, K., Kaiser, R. and Hoffmann, D., 2001, Geno2pheno: interpreting genotypic HIV drug resistance tests. IEEE Intelligent Systems 16, 35–41.

Berkelaar, M., Eikland, K. and Notebaert, P. 2004. Lp solve: open source (mixed-integer) linear programming system.

Caskey, M., Klein, F., Lorenzi, J.C., Seaman, M.S., West, A.P., Jr., Buckley, N., Kremer, G., Nogueira, L., Braunschweig, M., Scheid, J.F., Horwitz, J.A., Shimeliovich, I., Ben-Avraham, S., Witmer-Pack, M., Platten, M., Lehmann, C., Burke, L.A., Hawthorne, T., Gorelick, R.J., Walker, B.D., Keler, T., Gulick, R.M., Fatkenheuer, G., Schlesinger, S.J. and Nussenzweig, M.C., 2015, Viraemia suppressed in HIV-1-infected humans by broadly neutralizing antibody 3BNC117. Nature 522, 487–91.

Caskey, M., Schoofs, T., Gruell, H., Settler, A., Karagounis, T., Kreider, E.F., Murrell, B., Pfeifer, N., Nogueira, L., Oliveira, T.Y., Learn, G.H., Cohen, Y.Z., Lehmann, C., Gillor, D., Shimeliovich, I., Unson-O’Brien, C., Weiland, D., Robles, A., Kummerle, T., Wyen, C., Levin, R., Witmer-Pack, M., Eren, K., Ignacio, C., Kiss, S., West, A.P., Jr., Mouquet, H., Zingman, B.S., Gulick, R.M., Keler, T., Bjorkman, P.J., Seaman, M.S., Hahn, B.H., Fatkenheuer, G., Schlesinger, S.J., Nussenzweig, M.C. and Klein, F., 2017, Antibody 10-1074 suppresses viremia in HIV-1-infected individuals. Nat Med 23, 185–191.

Chuang, L.Y., Cheng, Y.H. and Yang, C.H., 2012, URPD: a specific product primer design tool. BMC Res Notes 5, 306.

DeKosky, B.J., Ippolito, G.C., Deschner, R.P., Lavinder, J.J., Wine, Y., Rawlings, B.M., Varadarajan, N., Giesecke, C., Dorner, T., Andrews, S.F., Wilson, P.C., Hunicke-Smith, S.P., Willson, C.G., Ellington, A.D. and Georgiou, G., 2013, High-throughput sequencing of the paired human immunoglobulin heavy and light chain repertoire. Nat Biotechnol 31, 166–9.

Doring, M., Buch, J., Friedrich, G., Pironti, A., Kalaghatgi, P., Knops, E., Heger, E., Obermeier, M., Daumer, M., Thielen, A., Kaiser, R., Lengauer, T. and Pfeifer, N., 2018, geno2pheno[ngs-freq]: a genotypic interpretation system for identifying viral drug resistance using next-generation sequencing data. Nucleic Acids Res 46, W271–W277.

Doring, M., Kreer, C., Lehnen, N., Klein, F. and Pfeifer, N., 2019, Modeling the Amplification of Immunoglobulins through Machine Learning on Sequence-Specific Features. Sci Rep 9, 10748.

Emrich, S.J., Lowe, M. and Delcher, A.L., 2003, PROBEmer: A web-based software tool for selecting optimal DNA oligos. Nucleic Acids Res 31, 3746–50.

Feige, U., 1998, A threshold of ln *n* for approximating set cover. J. ACM 45, 634–652.

Freund, N.T., Scheid, J.F., Mouquet, H. and Nussenzweig, M.C., 2015, Amplification of highly mutated human Ig lambda light chains from an HIV-1 infected patient. J Immunol Methods 418, 61–5.

Galson, J.D., Clutterbuck, E.A., Truck, J., Ramasamy, M.N., Munz, M., Fowler, A., Cerundolo, V., Pollard, A.J., Lunter, G. and Kelly, D.F., 2015, BCR repertoire sequencing: different patterns of B-cell activation after two Meningococcal vaccines. Immunol Cell Biol 93, 885–95.

Gardner, S.N., Hiddessen, A.L., Williams, P.L., Hara, C., Wagner, M.C. and Colston, B.W., Jr., 2009, Multiplex primer prediction software for divergent targets. Nucleic Acids Res 37, 6291–304.

Gardner, S.N., Jaing, C.J., Elsheikh, M.M., Pena, J., Hysom, D.A. and Borucki, M.K., 2014, Multiplex degenerate primer design for targeted whole genome amplification of many viral genomes. Adv Bioinformatics 2014, 101894.

Gautam, R., Nishimura, Y., Gaughan, N., Gazumyan, A., Schoofs, T., Buckler-White, A., Seaman, M.S., Swihart, B.J., Follmann, D.A., Nussenzweig, M.C. and Martin, M.A., 2018, A single injection of crystallizable fragment domain-modified antibodies elicits durable protection from SHIV infection. Nat Med 24, 610–616.

Giegerich, R., Meyer, F. and Schleiermacher, C., 1996, GeneFisher--software support for the detection of postulated genes. Proc Int Conf Intell Syst Mol Biol 4, 68–77.

Gupta, N.T., Vander Heiden, J.A., Uduman, M., Gadala-Maria, D., Yaari, G. and Kleinstein, S.H., 2015, Change-O: a toolkit for analyzing large-scale B cell immunoglobulin repertoire sequencing data. Bioinformatics 31, 3356–8.

Huang, Y.C., Chang, C.F., Chan, C.H., Yeh, T.J., Chang, Y.C., Chen, C.C. and Kao, C.Y., 2005, Integrated minimum-set primers and unique probe design algorithms for differential detection on symptom-related pathogens. Bioinformatics 21, 4330–7.

Hysom, D.A., Naraghi-Arani, P., Elsheikh, M., Carrillo, A.C., Williams, P.L. and Gardner, S.N., 2012, Skip the alignment: degenerate, multiplex primer and probe design using K-mer matching instead of alignments. PLoS One 7, e34560.

Ippolito, G.C., Hoi, K.H., Reddy, S.T., Carroll, S.M., Ge, X., Rogosch, T., Zemlin, M., Shultz, L.D., Ellington, A.D., Vandenberg, C.L. and Georgiou, G., 2012, Antibody repertoires in humanized NOD-scid-IL2Rgamma(null) mice and human B cells reveals human-like diversification and tolerance checkpoints in the mouse. PLoS One 7, e35497.

Jabado, O.J., Palacios, G., Kapoor, V., Hui, J., Renwick, N., Zhai, J., Briese, T. and Lipkin, W.I., 2006, Greene SCPrimer: a rapid comprehensive tool for designing degenerate primers from multiple sequence alignments. Nucleic Acids Res 34, 6605–11.

Jarman, S.N., 2004, Amplicon: software for designing PCR primers on aligned DNA sequences. Bioinformatics 20, 1644–5.

Kalendar, R., Lee, D. and Schulman, A.H., 2014, FastPCR software for PCR, in silico PCR, and oligonucleotide assembly and analysis. Methods Mol Biol 1116, 271–302.

Kampke, T., Kieninger, M. and Mecklenburg, M., 2001, Efficient primer design algorithms. Bioinformatics 17, 214–25.

Kaplon, H. and Reichert, J.M., 2018, Antibodies to watch in 2018. MAbs 10, 183–203.

Klein, F., Gaebler, C., Mouquet, H., Sather, D.N., Lehmann, C., Scheid, J.F., Kraft, Z., Liu, Y., Pietzsch, J., Hurley, A., Poignard, P., Feizi, T., Morris, L., Walker, B.D., Fatkenheuer, G., Seaman, M.S., Stamatatos, L. and Nussenzweig, M.C., 2012, Broad neutralization by a combination of antibodies recognizing the CD4 binding site and a new conformational epitope on the HIV-1 envelope protein. J Exp Med 209, 1469–79.

Klein, F., Mouquet, H., Dosenovic, P., Scheid, J.F., Scharf, L. and Nussenzweig, M.C., 2013, Antibodies in HIV-1 vaccine development and therapy. Science 341, 1199–204.

Kuiken, C., Korber, B. and Shafer, R.W., 2003, HIV sequence databases. AIDS Rev 5, 52–61.

Kuppers, R., Zhao, M., Hansmann, M.L. and Rajewsky, K., 1993, Tracing B cell development in human germinal centres by molecular analysis of single cells picked from histological sections. EMBO J 12, 4955–67.

Le Novere, N., 2001, MELTING, computing the melting temperature of nucleic acid duplex. Bioinformatics 17, 1226–7.

Lee, C., Wu, J.S., Shiue, Y.L. and Liang, H.L., 2006, MultiPrimer: software for multiplex primer design. Appl Bioinformatics 5, 99–109.

Lefranc, M.P., Giudicelli, V., Ginestoux, C., Bodmer, J., Muller, W., Bontrop, R., Lemaitre, M., Malik, A., Barbie, V. and Chaume, D., 1999, IMGT, the international ImMunoGeneTics database. Nucleic Acids Res 27, 209–12.

Lim, T.S., Mollova, S., Rubelt, F., Sievert, V., Dubel, S., Lehrach, H. and Konthur, Z., 2010, V-gene amplification revisited - An optimised procedure for amplification of rearranged human antibody genes of different isotypes. N Biotechnol 27, 108–17.

Linhart, C. and Shamir, R., 2002, The degenerate primer design problem. Bioinformatics 18 Suppl 1, S172–81.

Lorenz, R., Bernhart, S.H., Honer Zu Siederdissen, C., Tafer, H., Flamm, C., Stadler, P.F. and Hofacker, I.L., 2011, ViennaRNA Package 2.0. Algorithms Mol Biol 6, 26.

Markham, N.R. and Zuker, M., 2008, UNAFold: software for nucleic acid folding and hybridization. Methods Mol Biol 453, 3–31.

Mendoza, P., Gruell, H., Nogueira, L., Pai, J.A., Butler, A.L., Millard, K., Lehmann, C., Suarez, I., Oliveira, T.Y., Lorenzi, J.C.C., Cohen, Y.Z., Wyen, C., Kummerle, T., Karagounis, T., Lu, C.L., Handl, L., Unson-O’Brien, C., Patel, R., Ruping, C., Schlotz, M., Witmer-Pack, M., Shimeliovich, I., Kremer, G., Thomas, E., Seaton, K.E., Horowitz, J., West, A.P., Jr., Bjorkman, P.J., Tomaras, G.D., Gulick, R.M., Pfeifer, N., Fatkenheuer, G., Seaman, M.S., Klein, F., Caskey, M. and Nussenzweig, M.C., 2018, Combination therapy with anti-HIV-1 antibodies maintains viral suppression. Nature 561, 479–484.

Muellenbeck, M.F., Ueberheide, B., Amulic, B., Epp, A., Fenyo, D., Busse, C.E., Esen, M., Theisen, M., Mordmuller, B. and Wardemann, H., 2013, Atypical and classical memory B cells produce Plasmodium falciparum neutralizing antibodies. J Exp Med 210, 389–99.

Murugan, R., Imkeller, K., Busse, C.E. and Wardemann, H., 2015, Direct high-throughput amplification and sequencing of immunoglobulin genes from single human B cells. Eur J Immunol 45, 2698–700.

O’Halloran, D.M., 2016, PrimerMapper: high throughput primer design and graphical assembly for PCR and SNP detection. Sci Rep 6, 20631.

Ozawa, T., Kishi, H. and Muraguchi, A., 2006, Amplification and analysis of cDNA generated from a single cell by 5’-RACE: application to isolation of antibody heavy and light chain variable gene sequences from single B cells. Biotechniques 40, 469–70, 472, 474 passim.

Pappas, L., Foglierini, M., Piccoli, L., Kallewaard, N.L., Turrini, F., Silacci, C., Fernandez-Rodriguez, B., Agatic, G., Giacchetto-Sasselli, I., Pellicciotta, G., Sallusto, F., Zhu, Q., Vicenzi, E., Corti, D. and Lanzavecchia, A., 2014, Rapid development of broadly influenza neutralizing antibodies through redundant mutations. Nature 516, 418–22.

Pesole, G., Liuni, S., Grillo, G., Belichard, P., Trenkle, T., Welsh, J. and McClelland, M., 1998, GeneUp: a program to select short PCR primer pairs that occur in multiple members of sequence lists. Biotechniques 25, 112-7, 120–3.

Rachlin, J., Ding, C., Cantor, C. and Kasif, S., 2005, MuPlex: multi-objective multiplex PCR assay design. Nucleic Acids Res 33, W544–7.

Rollenske, T., Szijarto, V., Lukasiewicz, J., Guachalla, L.M., Stojkovic, K., Hartl, K., Stulik, L., Kocher, S., Lasitschka, F., Al-Saeedi, M., Schroder-Braunstein, J., von Frankenberg, M., Gaebelein, G., Hoffmann, P., Klein, S., Heeg, K., Nagy, E., Nagy, G. and Wardemann, H., 2018, Cross-specificity of protective human antibodies against Klebsiella pneumoniae LPS O-antigen. Nat Immunol 19, 617–624.

Rose, T.M., Henikoff, J.G. and Henikoff, S., 2003, CODEHOP (COnsensus-DEgenerate Hybrid Oligonucleotide Primer) PCR primer design. Nucleic Acids Res 31, 3763–6.

Rose, T.M., Schultz, E.R., Henikoff, J.G., Pietrokovski, S., McCallum, C.M. and Henikoff, S., 1998, Consensus-degenerate hybrid oligonucleotide primers for amplification of distantly related sequences. Nucleic Acids Res 26, 1628–35.

Rychlik, W., 2007, OLIGO 7 primer analysis software. Methods Mol Biol 402, 35–60.

Sblattero, D. and Bradbury, A., 1998, A definitive set of oligonucleotide primers for amplifying human V regions. Immunotechnology 3, 271–8.

Scheid, J.F., Mouquet, H., Ueberheide, B., Diskin, R., Klein, F., Oliveira, T.Y., Pietzsch, J., Fenyo, D., Abadir, A., Velinzon, K., Hurley, A., Myung, S., Boulad, F., Poignard, P., Burton, D.R., Pereyra, F., Ho, D.D., Walker, B.D., Seaman, M.S., Bjorkman, P.J., Chait, B.T. and Nussenzweig, M.C., 2011, Sequence and structural convergence of broad and potent HIV antibodies that mimic CD4 binding. Science 333, 1633–7.

Schneider, T.D., 2002, Consensus sequence Zen. Appl Bioinformatics 1, 111–9.

Shen, Z., Qu, W., Wang, W., Lu, Y., Wu, Y., Li, Z., Hang, X., Wang, X., Zhao, D. and Zhang, C., 2010, MPprimer: a program for reliable multiplex PCR primer design. BMC Bioinformatics 11, 143.

Sliepen, K., van Montfort, T., Ozorowski, G., Pritchard, L.K., Crispin, M., Ward, A.B. and Sanders, R.W., 2015, Engineering and Characterization of a Fluorescent Native-Like HIV-1 Envelope Glycoprotein Trimer. Biomolecules 5, 2919–34.

Souvenir, R., Buhler, J., Stormo, G. and Zhang, W. 2003 Selecting Degenerate Multiplex PCR Primers. In: G. Benson and R.D.M. Page (Eds.) Algorithms in Bioinformatics. Springer Berlin Heidelberg, Berlin, Heidelberg, p. 512–526.

Srivastava, G.P. and Xu, D., 2007, Genome-scale probe and primer design with PRIMEGENS. Methods Mol Biol 402, 159–76.

Sun, Y., Liu, H.Y., Mu, L. and Luo, E.J., 2012, Degenerate primer design to clone the human repertoire of immunoglobulin heavy chain variable regions. World J Microbiol Biotechnol 28, 381–6.

Tan, Y.G., Wang, Y.Q., Zhang, M., Han, Y.X., Huang, C.Y., Zhang, H.P., Li, Z.M., Wu, X.L., Wang, X.F., Dong, Y., Zhu, H.M., Zhu, S.D., Li, H.M., Li, N., Yan, H.P. and Gao, Z.H., 2016, Clonal Characteristics of Circulating B Lymphocyte Repertoire in Primary Biliary Cholangitis. J Immunol 197, 1609–20.

Taylor, B.S., Sobieszczyk, M.E., McCutchan, F.E. and Hammer, S.M., 2008, The challenge of HIV-1 subtype diversity. N Engl J Med 358, 1590–602.

Tiller, T., Meffre, E., Yurasov, S., Tsuiji, M., Nussenzweig, M.C. and Wardemann, H., 2008, Efficient generation of monoclonal antibodies from single human B cells by single cell RT-PCR and expression vector cloning. J Immunol Methods 329, 112–24.

Vander Heiden, J.A., Yaari, G., Uduman, M., Stern, J.N., O’Connor, K.C., Hafler, D.A., Vigneault, F. and Kleinstein, S.H., 2014, pRESTO: a toolkit for processing high-throughput sequencing raw reads of lymphocyte receptor repertoires. Bioinformatics 30, 1930–2.

Wang, J., Li, K.B. and Sung, W.K., 2004, G-PRIMER: greedy algorithm for selecting minimal primer set. Bioinformatics 20, 2473–5.

Wardemann, H., Yurasov, S., Schaefer, A., Young, J.W., Meffre, E. and Nussenzweig, M.C., 2003, Predominant autoantibody production by early human B cell precursors. Science 301, 1374–7.

Wright, E.S., Yilmaz, L.S., Ram, S., Gasser, J.M., Harrington, G.W. and Noguera, D.R., 2014, Exploiting extension bias in polymerase chain reaction to improve primer specificity in ensembles of nearly identical DNA templates. Environ Microbiol 16, 1354–65.

Wu, Y.C., Kipling, D., Leong, H.S., Martin, V., Ademokun, A.A. and Dunn-Walters, D.K., 2010, High-throughput immunoglobulin repertoire analysis distinguishes between human IgM memory and switched memory B-cell populations. Blood 116, 1070–8.

Yamada, T., Soma, H. and Morishita, S., 2006, PrimerStation: a highly specific multiplex genomic PCR primer design server for the human genome. Nucleic Acids Res 34, W665–9.

Ye, J., Ma, N., Madden, T.L. and Ostell, J.M., 2013, IgBLAST: an immunoglobulin variable domain sequence analysis tool. Nucleic Acids Res 41, W34–40.

Zhou, T., Georgiev, I., Wu, X., Yang, Z.Y., Dai, K., Finzi, A., Kwon, Y.D., Scheid, J.F., Shi, W., Xu, L., Yang, Y., Zhu, J., Nussenzweig, M.C., Sodroski, J., Shapiro, L., Nabel, G.J., Mascola, J.R. and Kwong, P.D., 2010, Structural basis for broad and potent neutralization of HIV-1 by antibody VRC01. Science 329, 811–7.

