## Supplementary Information for "openPrimeR for multiplex amplification of highly diverse templates"

#### Supplementary Figures

| | |
|---|---|
| Figure S1 | *openPrimeR graphical user interface* |
| Figure S2 | *Visualization of PCR products for IGHV, IGKV, and IGLV gene libraries by gelelectrophoresis* |
| Figure S3 | *Gating strategies for single cell sorting* |
| Figure S4 | *Visualization of PCR products from naive B cell cDNA with oPR-IGHV, Set 1, and Set 2* |
| Figure S5 | *Visualization of PCR products from IgG+ memory B cell cDNA with oPR-IGHV, Set 1, and Set 2* |
| Figure S6 | *Visualization of PCR products from HIV-1$_{BG505.SOSIP}$-reactive B cell cDNA with oPR-IGHV, Set 1, and Set 2* |
| Figure S7 | *PCR results from HIV-1$_{BG505.SOSIP}$-reactive B cell cDNA with oPR-IGKV and oPR-IGLV* |

#### Supplementary Tables

| | |
|---|---|
| Table S1 | *openPrimeR options, default settings and settings used for primer design on IGHV, IGKV, and IGLV* |
| Table S2 | *IGH, IGK, and IGL V gene alleles used in standard libraries* |
| Table S3 | *openPrimeR-derived optimized primer sets oPR-IGHV, oPR-IGKV, and oPR-IGLV for amplification of antibody heavy and light chains* |
| Table S4 | *Primer combinations and thermocycling conditions for multiplex PCRs* |

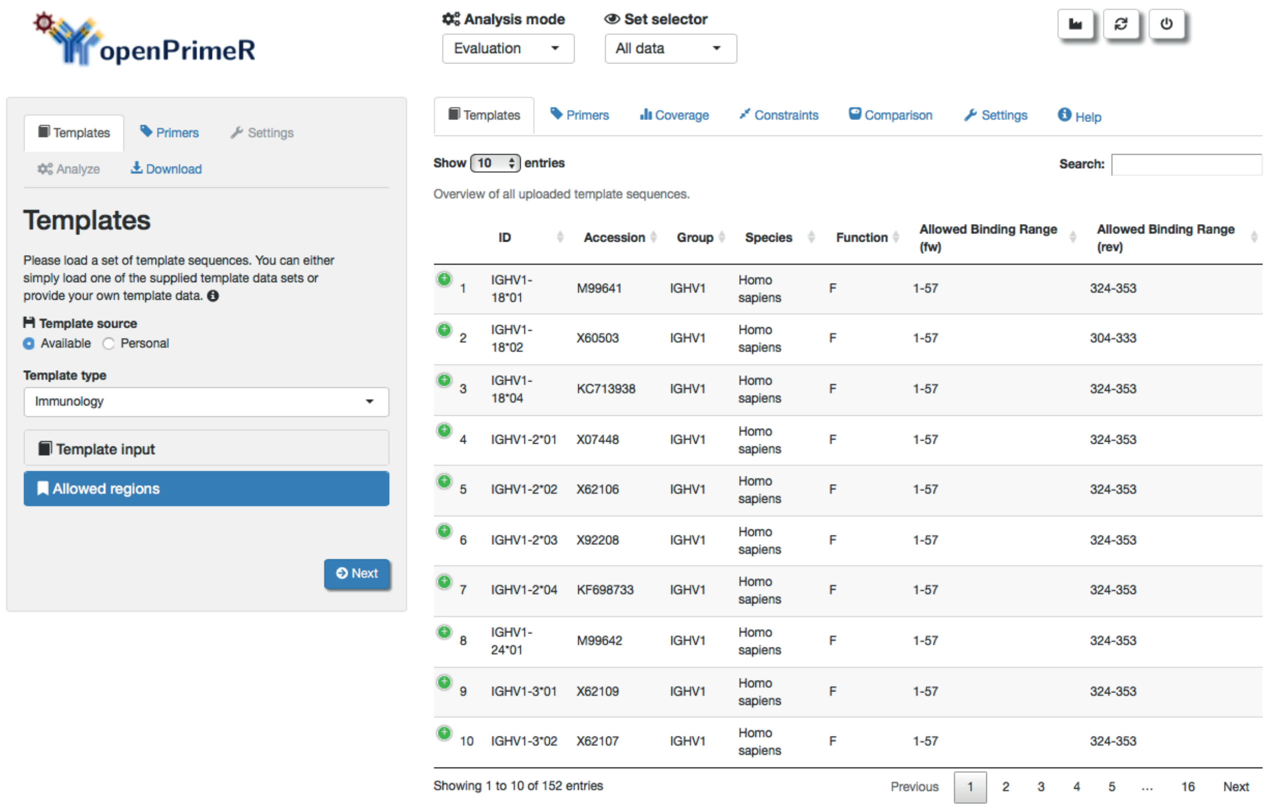

**Figure S1: openPrimeR graphical user interface.** A Shiny application can be used for interactive usage of openPrimeR. The left panel contains the input interfaces and guides the user step-wise through the evaluation or design mode. The right panel offers several tabs for output display.

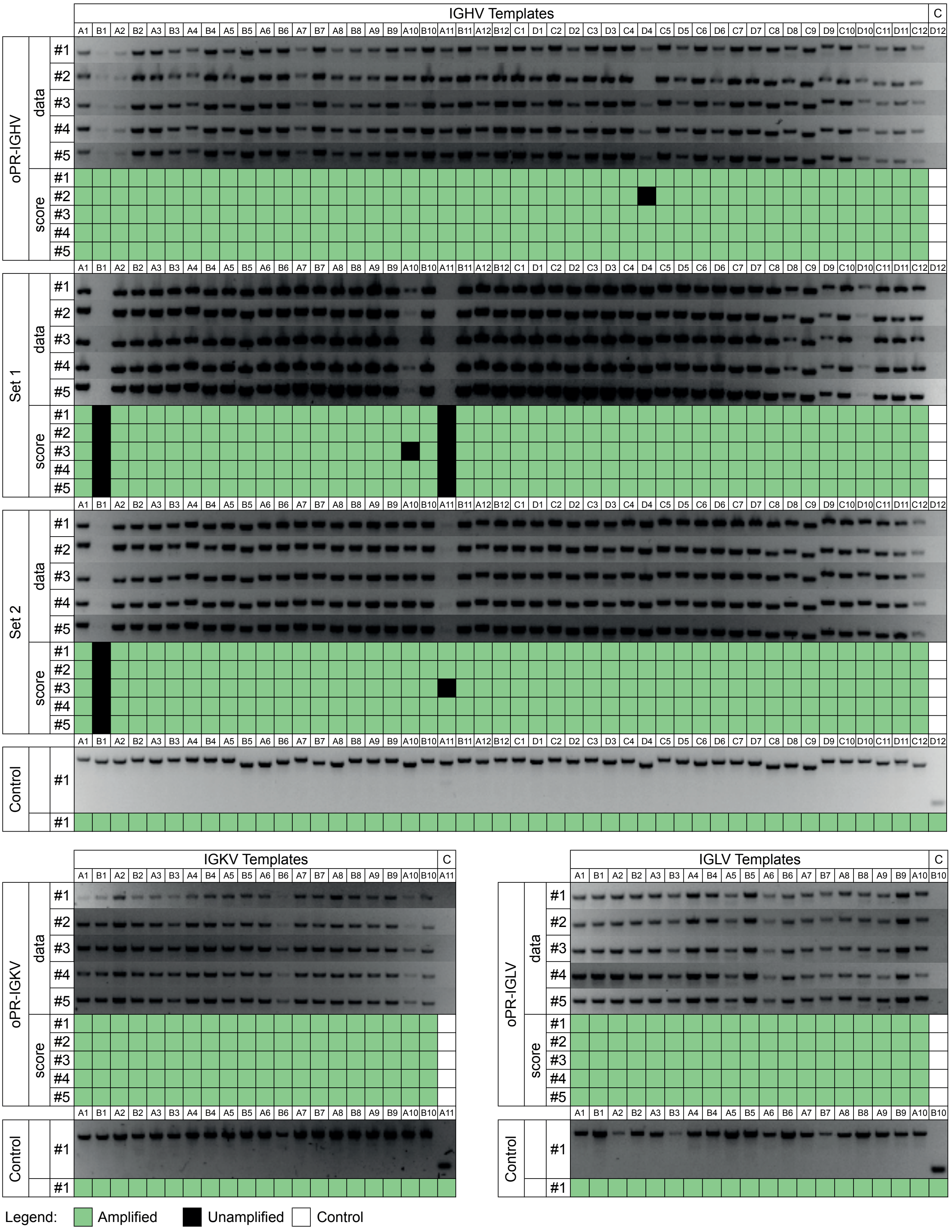

**Figure S2: Visualization of PCR products for IGHV, IGKV, and IGLV gene libraries by gel-electrophoresis.** IGHV library was amplified with oPR-IGHV, Set 1, and Set 2. IGKV and IGLV gene libraries were amplified with oPR-IGKV and oPR-IGLV, respectively. Green, black, and white squares indicate amplified, unamplified, and control wells (not counted). Note: IGHV template in D4 turned out to have a deletion in the binding region of oPR-IGHV primer 9.

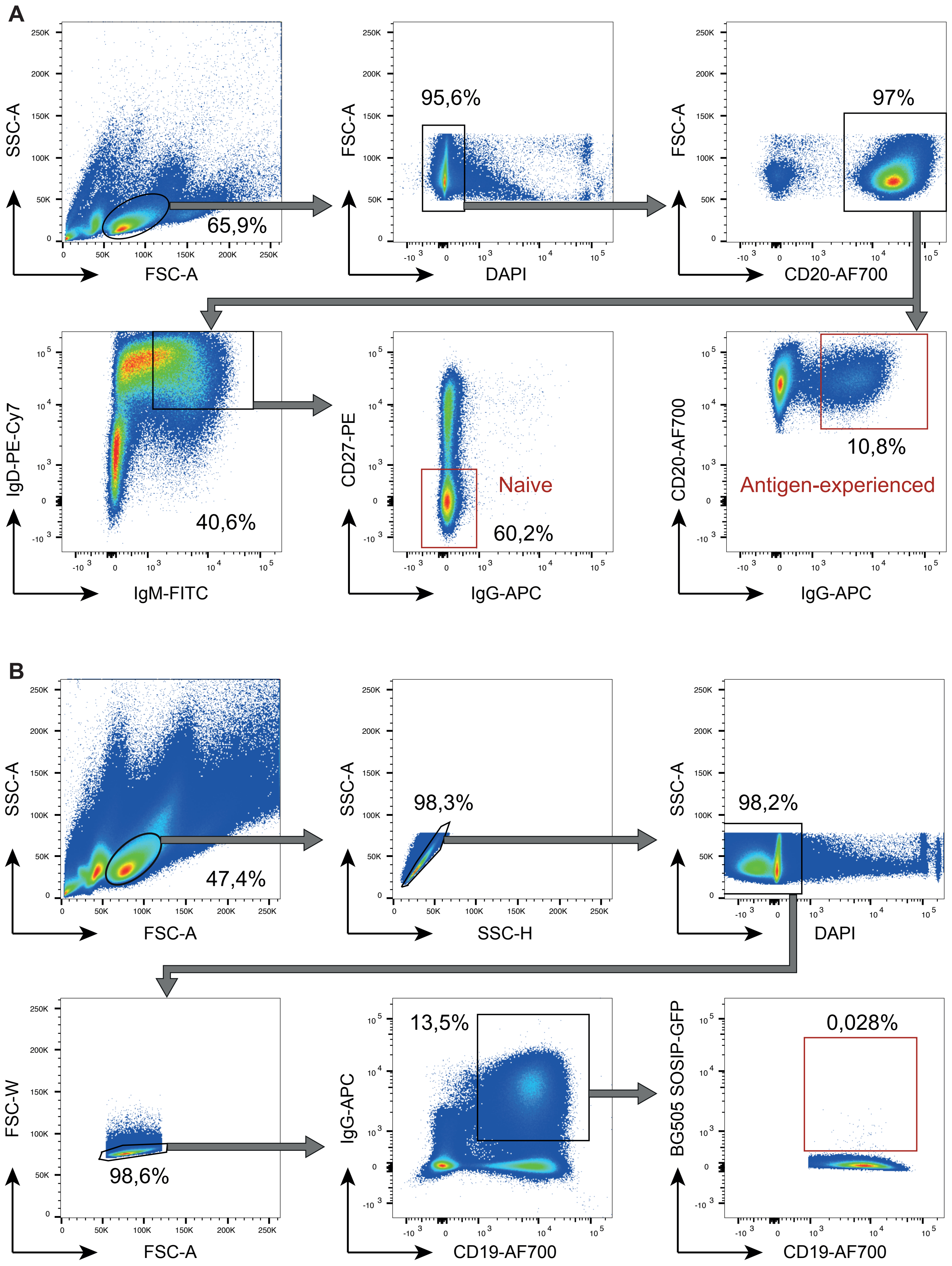

**Figure S3: Gating strategies for single cell sorting.** (**A**) Gating strategy for naive and antigen-experienced B cells. (**B**) Gating strategy for HIV-1-specific B cells.

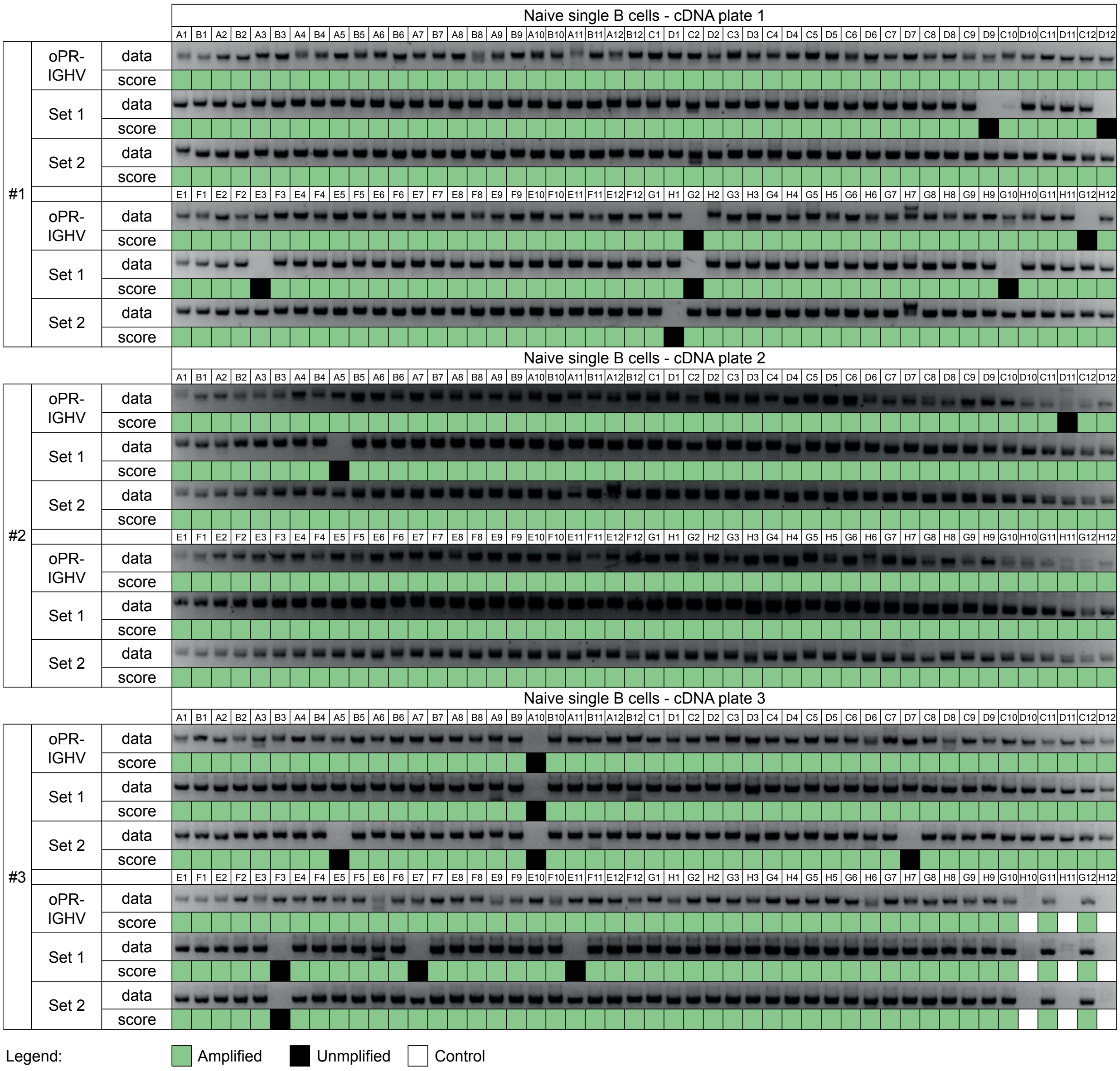

**Figure S4: Visualization of PCR products from naive B cell cDNA with oPR-IGHV, Set 1, and Set 2.** Green, black, and white squares indicate amplified, unamplified, and control wells (not counted), respectively.

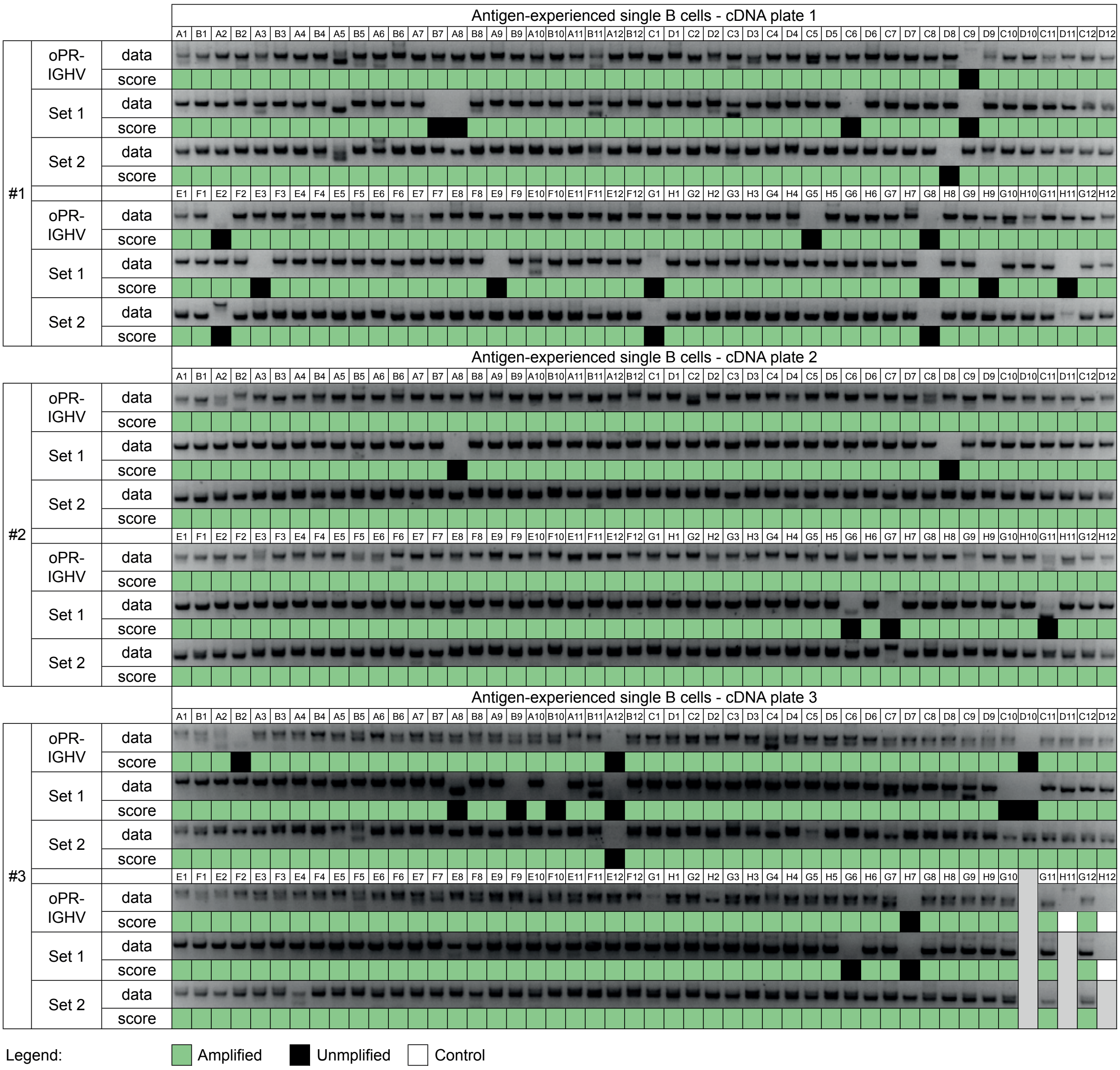

**Figure S5: Visualization of PCR products from IgG⁺ memory B cell cDNA with oPR-IGHV, Set 1, and Set 2.** Green, black, and white squares indicate amplified, unamplified, and control wells, respectively. Note: Controls and grey squares were not taken into account for coverage determination.

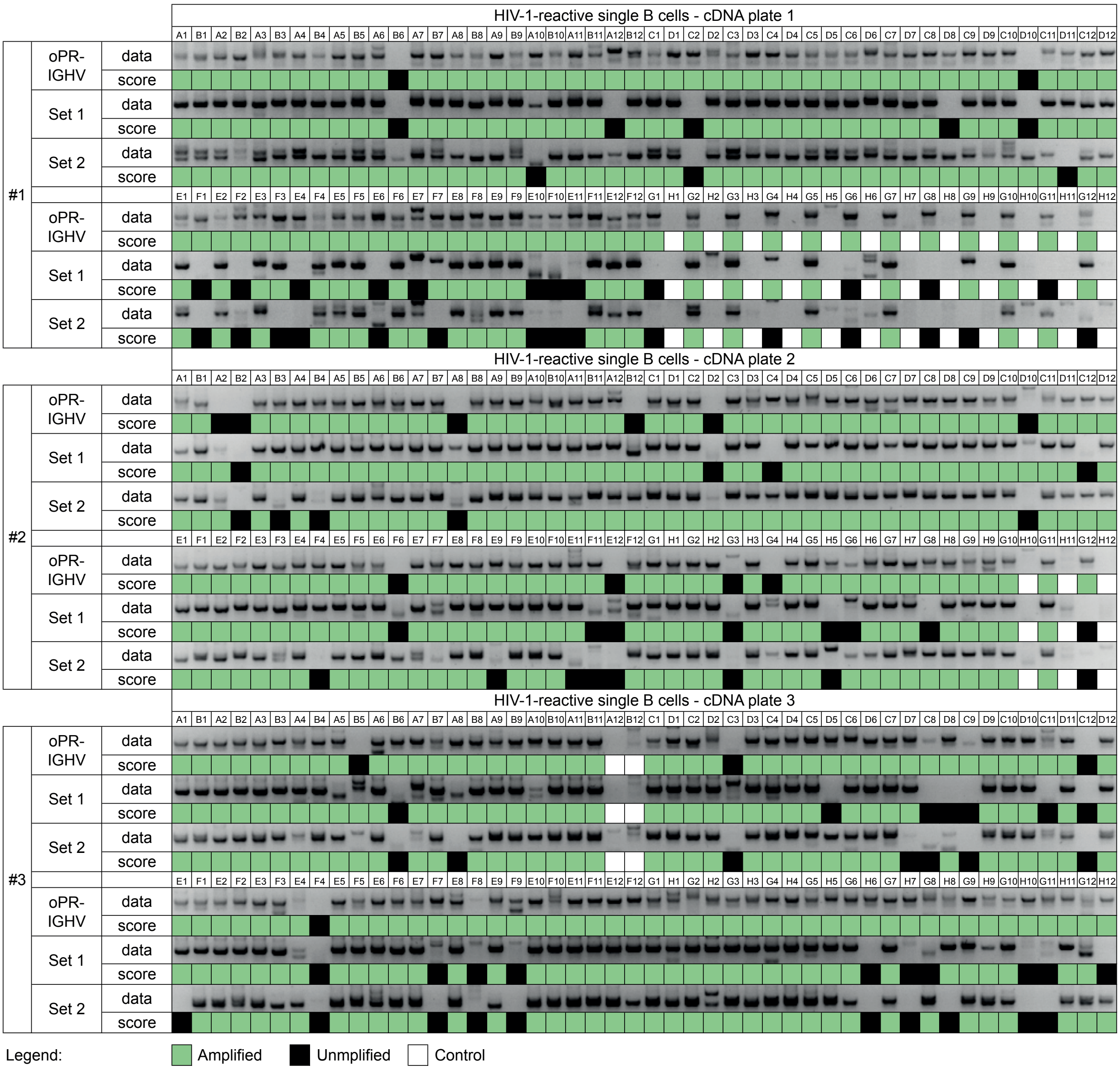

**Figure S6: Visualization of PCR products from HIV-1$_{BG505.SOSIP}$-reactive B cell cDNA with oPR-IGHV, Set 1, and Set 2.** Green, black, and white squares indicate amplified, unamplified, and control wells (not counted), respectively.

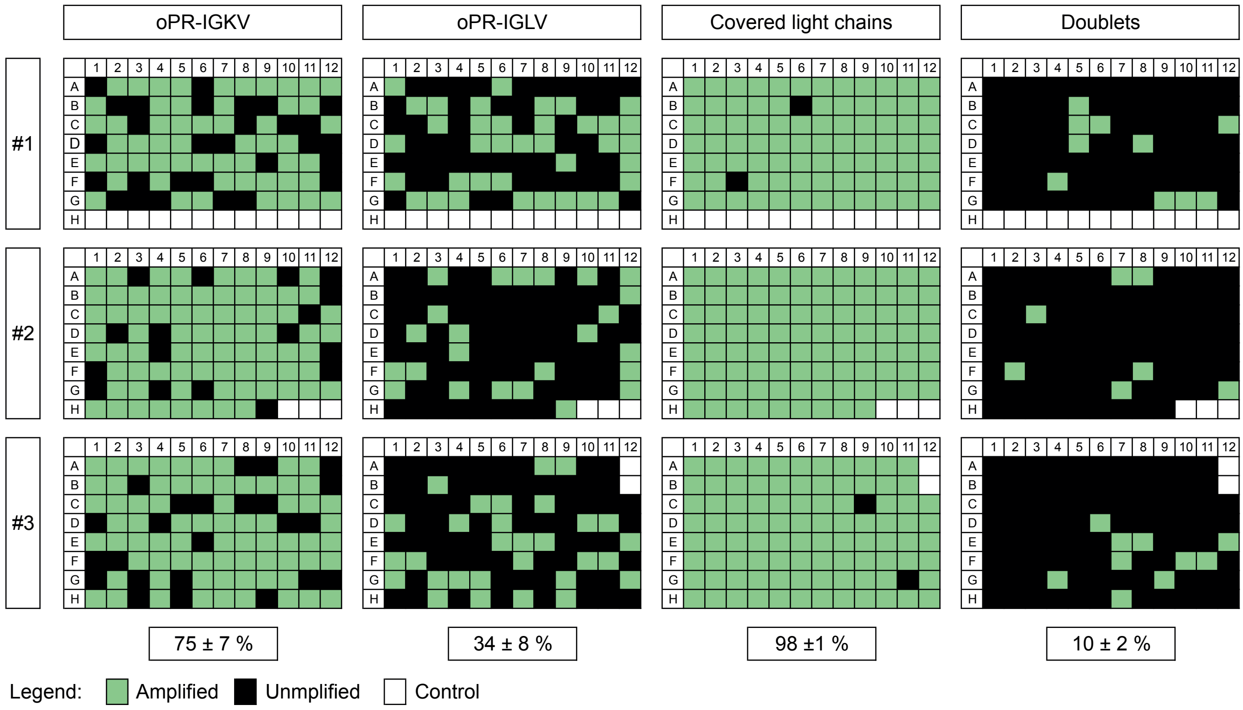

**Figure S7: PCR results from HIV-1$_{BG505.SOSIP}$-reactive B cell cDNA with oPR-IGKV and oPR-IGLV (related to Figure 6).** Light chain coverage summary. Kappa and lambda light chain specific PCRs were performed on HIV-1$_{BG505.SOSIP}$ reactive B cells with oPR-IGKV and oPR-IGLV primer sets. Green, black, and white boxes depict amplified, unamplified, and control wells (not counted), respectively. Percentages indicate mean coverage ± standard deviation.

**Table S1**. openPrimeR options, default settings and settings used for primer design on IGHV, IGKV, and IGLV.

| Options | Ranges | | Default values | |
|---|---|---|---|---|
| | Settings | Limits | Settings | Limits |
| **Template** | | | | |
| Template source | preloaded/personal | - | preloaded | - |
| Binding region | complete tempalte | - | complete leader | - |
| **Primers** | | | | |
| Template sequence relationship | related/divergent | - | related | - |
| Target strand for deisgn | Sense/antisense/both | - | both | - |
| Degeneracy | 1 to 64 | - | 1 | - |
| **Coverage conditions** | | | | |
| Allowed mismatches | 0 to 20 | - | 7 | - |
| **Binding conditions** | | | | |
| Annealing energy* | off/-20 to 0 kcal/mol | - | off | - |
| Amplification efficiency† | off/0.1 to 100 % | - | off | - |
| Coverage model‡ | off/0 to 100 % | - | 6% | - |
| Forbidden number of 3' Mismatches | 0 (allowed) to 10 | - | 0 (allowed) | - |
| **Codon design** | | | | |
| Introduction of stopcodons by mismatches | allowed/forbidden | - | allowed | - |
| Aminoacid substitutions by mismatch binding | allowed/forbidden | - | allowed | - |
| **Constraints** | | | | |
| Minimal number of individual targets | off/1 to 100 | - | 1 | - |
| Primer length | off/1 to 50 | - | 18-23 | - |
| GC clamp | off/0 to 10 | 0 to 5 | 1 to 3 | 0 to 4 |
| GC ratio | off/0 to 100% | 0 to 100% | 40 to 60% | 30 to 70% |
| Run length | off/0 to 10 | 0 to 20 | 0 to 4 | 0 to 6 |
| Repeat length | off/0 to 10 | 0 to 20 | 0 to 4 | 0 to 6 |
| Melting temperature range | off/40 to 90°C | 40 to 90°C | 50 to 70°C | 50 to 75°C |
| Melting temperature deviation | off/0 to 15°C | 0 to 20°C | 0 to 5°C | 0 to 8°C |
| Secondary structure ΔG# | off/-10 to 0 kcal/mol | -10 to 0 kcal/mol | -1 kcal/mol | -2 kcal/mol |
| Specificity | off/0 to 100% | 0 to 100% | 100% | 80 to 100% |
| Self-dimerization ΔG# | off/-15 to 0 kcal/mol | -20 to 0 kcal/mol | -5 kcal/mol | -7 kcal/mol |
| Cross-dimerization ΔG# | off/-15 to 0 kcal/mol | -20 to 0 kcal/mol | -5 kcal/mol | -7 kcal/mol |
| **PCR conditions** | | | | |
| Polymerase | Taq/Non-Taq | - | Taq | - |
| Number of PCR cycles | 1 to 100 | - | 25 | - |
| $Na^+$ concentration | 0 to 100 mM | - | 0 | - |
| $Mg^{2+}$ concentration | 0 to 100 mM | - | 1.5 | - |
| $K^+$ concentration | 0 to 100 mM | - | 50 | - |
| Tris concentration | 0 to 100 mM | - | 0 | - |
| Primer concentration | 0 to 1000 nM | - | 200 | - |
| Template concentration | 0.01 to 1000 nM | - | 0.01 | - |
| **Design options** | | | | |
| Optimization algorithm | Greedy/ILP | - | Greedy | - |
| Target coverage | 0 to 100 % | - | 100% | - |

* Annealing energy is determined with MELTING (Le Novere, 2001) and binding events are only counted if their binding energy is below the adjusted threshhold

† Amplification efficiency is determined with DECIPHER (Wright et al., 2014) and binding events are only counted if their efficiency is within the adjusted range

‡ A logistic regression model (Doring et al., accepted) estimates a false positive rate for each binding event prediction and only events below the adjusted threshhold are counted

### Free energy (ΔG) is calculated for secondary structures and dimerization events with OligoArrayAux (Markham and Zuker, 2008) and ViennaRNA (Lorenz et al., 2011), respectively. Primers with free energies below the adjusted threshhold are considered to violate the constraint

**Table S1**. openPrimeR options, default settings and settings used for primer design on IGHV, IGKV, and IGLV. (continued)

| Options | openPrimeR-IGHV | | openPrimeR-IGKV | | openPrimeR-IGLV | |
|---|---|---|---|---|---|---|
| | Settings | Limits | Settings | Limits | Settings | Limits |
| **Template** | | | | | | |
| Template source | preloaded | - | personal | - | personal | - |
| Binding region | -60 to -20 | - | -69 to -20 | - | -63 to -10 | - |
| **Primers** | | | | | | |
| Template sequence relationship | related | - | related | - | related | - |
| Target strand for deisgn | antisense | - | antisense | - | antisense | - |
| Degeneracy | 1 | - | 1 | - | 1 | - |
| **Coverage conditions** | | | | | | |
| Allowed mismatches | 1 | - | 1 | - | 1 | - |
| **Binding conditions** | | | | | | |
| Annealing energy* | off | - | off | - | off | - |
| Amplification efficiency† | off | - | off | - | off | - |
| Coverage model‡ | off | - | off | - | off | - |
| Forbidden number of 3' Mismatches | 1 | - | 3 | - | 3 | - |
| **Codon design** | | | | | | |
| Introduction of stopcodons by mismatches | forbidden | - | forbidden | - | forbidden | - |
| Aminoacid substitutions by mismatch binding | allowed | - | allowed | - | allowed | - |
| **Constraints** | | | | | | |
| Minimal number of individual targets | 1 | - | 1 | - | 1 | - |
| Primer length | 18-28 | - | 18-30 | - | 18-30 | - |
| GC clamp | 1 to 3 | 1 to 3 | 1 to 3 | 1 to 3 | 1 to 3 | 1 to 3 |
| GC ratio | 40 to 60% | 30 to 70% | 40 to 60% | 30 to 70% | 40 to 60% | 30 to 70% |
| Run length | 0 to 4 | 0 to 6 | 0 to 4 | 0 to 6 | 0 to 4 | 0 to 6 |
| Repeat length | 0 to 4 | 0 to 6 | 0 to 4 | 0 to 6 | 0 to 4 | 0 to 6 |
| Melting temperature range | 60 to 75°C | 57 to 78°C | 60 to 75°C | 57 to 78°C | 60 to 75°C | 57 to 78°C |
| Melting temperature deviation | 0 to 3°C | 0 to 3 °C | 0 to 3°C | 0 to 3 °C | 0 to 3°C | 0 to 3 °C |
| Secondary structure ΔG# | off | off | off | off | off | off |
| Specificity | 1 | 1 | 1 | 1 | 1 | 1 |
| Self-dimerization ΔG# | -5 kcal/mol | -5 kcal/mol | -5 kcal/mol | -5 kcal/mol | -5 kcal/mol | -5 kcal/mol |
| Cross-dimerization ΔG# | -5 kcal/mol | -5 kcal/mol | -5 kcal/mol | -5 kcal/mol | -5 kcal/mol | -5 kcal/mol |
| **PCR conditions** | | | | | | |
| Polymerase | Taq | - | Taq | - | Taq | - |
| Number of PCR cycles | 25 | - | 25 | - | 25 | - |
| $Na^+$ concentration | 0 | - | 0 | - | 0 | - |
| $Mg^{2+}$ concentration | 1.5 | - | 1.5 | - | 1.5 | - |
| $K^+$ concentration | 50 | - | 50 | - | 50 | - |
| Tris concentration | 0 | - | 0 | - | 0 | - |
| Primer concentration | 200 | - | 200 | - | 200 | - |
| Template concentration | 0.01 | - | 0.01 | - | 0.01 | - |
| **Design options** | | | | | | |
| Optimization algorithm | ILP | - | ILP | - | ILP | - |
| Target coverage | 1 | - | 1 | - | 1 | - |

* Annealing energy is determined with MELTING (Le Novere, 2001) and binding events are only counted if their binding energy is below the adjusted threshhold

† Amplification efficiency is determined with DECIPHER (Wright et al., 2014) and binding events are only counted if their efficiency is within the adjusted range

‡ A logistic regression model (Doring et al., accepted) estimates a false positive rate for each binding event prediction and only events below the adjusted threshhold are counted

### Free energy (ΔG) is calculated for secondary structures and dimerization events with OligoArrayAux (Markham and Zuker, 2008) and ViennaRNA (Lorenz et al., 2011), respectively. Primers with free energies below the adjusted threshhold are considered to violate the constraint

**Table S2**. IGH, IGK, and IGL V gene alleles used in standard libraries.

| IGHV standard library | | | | | | | | | | | |
|---|---|---|---|---|---|---|---|---|---|---|---|
| Well† | V gene | Allele | Well† | V gene | Allele | Well† | V gene | Allele | Well† | V gene | Allele |
| A1 | IGHV1-2 | 02 | B1 | IGHV2-70 | 15 | C1 | IGHV3-43 | 02 | D1 | IGHV4-30-2 | 01 |
| A2 | IGHV1-3 | 01 | B2 | IGHV3-7 | 01 | C2 | IGHV3-48 | 01 | D2 | IGHV4-31 | 03 |
| A3 | IGHV1-8 | 01 | B3 | IGHV3-9 | 01 | C3 | IGHV3-49 | 05 | D3 | IGHV4-34 | 01 |
| A4 | IGHV1-18 | 04 | B4 | IGHV3-11 | 01 | C4 | IGHV3-53 | 02 | D4 | IGHV4-38-2 | 02 |
| A5 | IGHV1-24 | 01 | B5 | IGHV3-13 | 01 | C5 | IGHV3-64 | 01 | D5 | IGHV4-39 | 01 |
| A6 | IGHV1-45 | 02 | B6 | IGHV3-15 | 01 | C6 | IGHV3-64D | 06 | D6 | IGHV4-59 | 01 |
| A7 | IGHV1-46 | 01 | B7 | IGHV3-20 | 01 | C7 | IGHV3-66 | 01 | D7 | IGHV4-61 | 01 |
| A8 | IGHV1-58 | 01 | B8 | IGHV3-21 | 01 | C8 | IGHV3-72 | 01 | D8 | IGHV5-10-1 | 03 |
| A9 | IGHV1-69 | 01 | B9 | IGHV3-23 | 01 | C9 | IGHV3-73 | 01 | D9 | IGHV5-51 | 03 |
| A10 | IGHV1-69-2 | 01 | B10 | IGHV3-30 | 18 | C10 | IGHV3-74 | 01 | D10 | IGHV6-1 | 01 |
| A11 | IGHV2-5 | 02 | B11 | IGHV3-30-3 | 01 | C11 | IGHV4-4 | 07 | D11 | IGHV7-4-1 | 02 |
| A12 | IGHV2-26 | 01 | B12 | IGHV3-33 | 01 | C12 | IGHV4-28 | 07 | D12 | | |

| IGKV standard library | | | | | | IGLV standard library | | | | | |
|---|---|---|---|---|---|---|---|---|---|---|---|
| Well† | V gene | Allele | Well† | V gene | Allele | Well† | V gene | Allele | Well† | V gene | Allele |
| A1 | IGKV1-5 | 01 | B1 | IGKV1-6 | 01 | A1 | IGLV1-40 | 01 | B1 | IGLV1-44 | 01 |
| A2 | IGKV1-8 | 01 | B2 | IGKV1-9 | 01 | A2 | IGLV1-47 | 01 | B2 | IGLV1-51 | 01 |
| A3 | IGKV1-12 | 01 | B3 | IGKV1-13 | 02 | A3 | IGLV2-11 | 01 | B3 | IGLV2-14 | 02 |
| A4 | IGKV1-16 | 02 | B4 | IGKV1-27 | 01 | A4 | IGLV2-18 | 02 | B4 | IGLV2-23 | 01 |
| A5 | IGKV1-39 | 01 | B5 | IGKV2-28 | 01 | A5 | IGLV3-1 | 01 | B5 | IGLV3-10 | 01 |
| A6 | IGKV2-29 | 01 | B6 | IGKV2-30 | 02 | A6 | IGLV3-19 | 01 | B6 | IGLV3-21 | 02 |
| A7 | IGKV2D-28 | 01 | B7 | IGKV2D-29 | 01 | A7 | IGLV3-25 | 02 | B7 | IGLV5-45 | 03 |
| A8 | IGKV3-11 | 01 | B8 | IGKV3-15 | 01 | A8 | IGLV4-60 | 03 | B8 | IGLV7-46 | 01 |
| A9 | IGKV3-20 | 01 | B9 | IGKV3D-15 | 01 | A9 | IGLV6-57 | 02 | B9 | IGLV10-54 | 01 |
| A10 | IGKV4-1 | 01 | B10 | IGKV6-21 | 01 | A10 | IGLV9-49 | 01 | B10 | | |
| A11 | | | B11 | | | A11 | | | B11 | | |
| A12 | | | B12 | | | A12 | | | B12 | | |

† Well notation indicates the position of the individual V gene allele on the standard library multiwell plate

**Table S3**. openPrimeR-derived optimized primer sets oPR-IGHV, oPR-IGKV, and oPR-IGLV for amplification of antibody heavy and light chains.

| **openPrimeR IGHV set (oPR-IGHV)** | | |
|---|---|---|
| **Name** | **Sequence** | **Length** |
| oPR-IGHV-1_fw | ATGGACTGGACCTGGAGCATCC | 22 |
| oPR-IGHV-2_fw | ATGGACTGGACCTGGAGGATCCTC | 24 |
| oPR-IGHV-3_fw | ATGGACTGGACCTGGAGGGTCTTC | 24 |
| oPR-IGHV-4_fw | ATGGACTGGATTTGGAGGGTCCTCTTC | 27 |
| oPR-IGHV-5_fw | ATGGACACACTTTGCTACACACTCCTGC | 28 |
| oPR-IGHV-6_fw | ACTTTGCTCCACGCTCCTGC | 20 |
| oPR-IGHV-7_fw | GGCTGAGCTGGGTTTTCCTTGTTG | 24 |
| oPR-IGHV-8_fw | GGCTCCGCTGGGTTTTCCTTGTTG | 24 |
| oPR-IGHV-9_fw | CACCTGTGGTTCTTCCTCCTGCTG | 24 |
| oPR-IGHV-10_fw | ATGAAACACCTGTGGTTCTTCCTCCTCC | 28 |
| oPR-IGHV-11_fw | ACATCTGTGGTTCTTCCTTCTCCTGGTG | 28 |
| oPR-IGHV-12_fw | GCCTCTCCACTTAAACCCAGGCTC | 24 |
| oPR-IGHV-13_fw | ATGTCTGTCTCCTTCCTCATCTTCCTGC | 28 |
| oPR-IGHV-14_fw | ATGGAGTTGGGGCTGAGCTGG | 21 |
| oPR-IGHV-15_fw | ATGGGGTCAACCGCCATCCTC | 21 |
| **openPrimeR IGKV set (oPR-IGKV)** | | |
| **Name** | **Sequence** | |
| oPR-IGKV-1_fw | ATGAGGCTCCTTGCTCAGCTTCTGG | 25 |
| oPR-IGKV-2_fw | ATGGAAGCCCCAGCTCAGCTTC | 22 |
| oPR-IGKV-3_fw | CCCAGCTCAGCTTCTCTTCCTCCTG | 25 |
| oPR-IGKV-4_fw | TGGTGTTGCAGACCCAGGTCTTCATTTC | 28 |
| oPR-IGKV-5_fw | GTCCCAGGTTCACCTCCTCAGCTTC | 25 |
| oPR-IGKV-6_fw | GCCATCACAACTCATTGGGTTTCTGCTG | 28 |
| oPR-IGKV-7_fw | TCCCTGCTCAGCTCCTGGG | 19 |
| oPR-IGKV-8_fw | CCTGGGACTCCTGCTGCTCTG | 21 |
| **openPrimeR IGHV set (oPR-IGLV)** | | |
| **Name** | **Sequence** | |
| oPR-IGLV-1_fw | CCCTGGGTCATGCTCCTCCTGAAATC | 26 |
| oPR-IGLV-2_fw | CTCTGCTGCTCCTCACTCTCCTCAC | 25 |
| oPR-IGLV-3_fw | ATGGCATGGATCCCTCTCTTCCTCG | 25 |
| oPR-IGLV-4_fw | CCTCTCTGGCTCACTCTCCTCACTC | 25 |
| oPR-IGLV-5_fw | ACACTCCTGCTCCCACTCCTCAAC | 24 |
| oPR-IGLV-6_fw | ATGGCCTGGATCCCTCTACTTCTCC | 25 |
| oPR-IGLV-7_fw | ATGGCCTGGGTCTCCTTCTACC | 22 |
| oPR-IGLV-8_fw | ATGGCCTGGACTCCTCTCTTTCTGTTC | 27 |
| oPR-IGLV-9_fw | ATGGCCTGGATGATGCTTCTCCTC | 24 |
| oPR-IGLV-10_fw | GTCCCCTCTCTTCCTCACCCTCATC | 25 |
| oPR-IGLV-11_fw | CTCCTCGCTCACTGCACAGG | 20 |
| oPR-IGLV-12_fw | CCTCTCCTCCTCACCCTCCTC | 21 |
| oPR-IGLV-13_fw | CTCCTCCTCACCCTCCTCACTC | 22 |
| oPR-IGLV-14_fw | ATGGCCTGGACCCCTCTCC | 19 |
| oPR-IGLV-15_fw | ATGGCCTGGACCCCACTCC | 19 |

**Table S4**. Primer combinations and thermocycling conditions for multiplex PCRs.

| Subset | PCR step | Primer | Set 1 | Set 2 | oPR-IGHV | oPR-IGKV | oPR-IGLV |
|---|---|---|---|---|---|---|---|
| Naive | 1st PCR | Forward | Set 1 Mix 1 | Set 2 Mix | oPR-IGHV Mix | - | - |
| | | Reverse | Cm-RT | Cm-RT | Cm-RT | - | - |
| | 2nd PCR | Forward | Set 1 Mix 2 | Set 2 Mix | oPR-IGHV Mix | - | - |
| | | Reverse | All IgM reverse | All IgM reverse | All IgM reverse | - | - |
| $IgG^+$ memory & HIV-1 reactive | 1st PCR | Forward | Set 1 Mix 1 | Set 2 Mix | oPR-IGHV Mix | oPR-IGKV Mix | oPR-IGLV Mix |
| | | Reverse | 3' Cγ CH1 | Cg-RT | Cg-RT | 3' Cκ 543 | 3' Cλ |
| | 2nd PCR | Forward | Set 1 Mix 2 | Set 2 Mix | oPR-IGHV Mix | oPR-IGKV Mix | oPR-IGLV Mix |
| | | Reverse | 3' IgG (internal) | All IgG reverse | 3' IgG (internal) | 3' Cκ 494 | 3' XhoI Cλ |

| PCR | Step | | Temperature (°C) | Time (sec) | Repeats |
|---|---|---|---|---|---|
| 1st PCR | 1 | Initial denaturation | 94 | 120 | 1 |
| | 2 | Denaturation | 94 | 30 | 50 |
| | | Annealing | 57 | 30 | |
| | | Extension | 72 | 55 | |
| | 3 | Final extension | 72 | 300 | 1 |
| | 4 | Hold | 4 | - | - |
| 2nd PCR | 1 | Initial denaturation | 94 | 120 | 1 |
| | 2 | Denaturation | 94 | 30 | 50 |
| | | Annealing | 57 | 30 | |
| | | Extension | 72 | 45 | |
| | 3 | Final extension | 72 | 300 | 1 |
| | 4 | Hold | 4 | - | - |
